# Disordered peptides impair neutrophil bacterial clearance and enhance tissue damage in septic patients

**DOI:** 10.1101/2020.07.31.227017

**Authors:** Csaba I. Timár, Ferenc Kolonics, Viktor Berzsenyi, Eszter Tamáska, Anna Párkányi, Michael L. Merchant, Daniel W. Wilkey, Zsolt Iványi, Kenneth R. McLeish, Erzsébet Ligeti

## Abstract

Neutrophilic granulocytes are required for antimicrobial defense, but they can also be harmful to the host organism. The current study demonstrates that disordered peptides in the 3-12 kDa size range in the plasma of septic patients alter effector functions of neutrophils from healthy donors. Those peptides stimulated exocytosis, increased extracellular release of reactive oxygen species (ROS), decreased ROS production in phagosomes, and impaired elimination of ROS-sensitive bacteria. Both the concentration of peptides in patients’ plasma and their effects on healthy cells were proportional to the clinical status of the patients. Proteomic analysis and in silico modeling indicate that multiple proteases generate the toxic peptides, with the greatest number of peptides cleaved by neutrophil elastase. We propose that neutrophils participate in an amplification loop in which proteolytic peptides stimulate extracellular release of proteases, resulting in production of more peptides. The enhanced extracellular ROS release contributes to tissue damage, while reduced intracellular ROS generation impairs elimination of certain bacteria. Breaking of this vicious cycle may offer a potential target for intervention.

## Introduction

Sepsis is a common, life-threatening complication of infection leading to a dysregulated host response and subsequent multi-organ failure (1),(2–6). In spite of modern antibiotic and intensive care therapy, the mortality varies between 15% and 60%, depending on the severity of the cases and on the causative organism (7–9). Several lines of evidence support involvement of neutrophilic granulocytes (PMN), including i.) massive accumulation of PMN at the site of injury or infection, ii.) significant alterations in the number and function of circulating PMN, and iii.) depletion of PMN combined with adequate antibiotics treatment improves the outcome (3, 10, 11), suggesting a dual (defending and harmful) role of PMN in the pathogenesis. In spite of intensive study, the role of PMN in sepsis remains poorly understood. Previous reports describe impaired PMN migration, (12–14) altered expression of adhesion molecules, (15, 16) increased lifespan, (17–19) and either increased (16, 20–22) or decreased (23, 24) production of reactive oxygen species (ROS). These studies typically investigated one or very few neutrophil functions, while global bactericidal activity and correlation between functional and clinical status were rarely evaluated (15, 25).

The aim of the present study was to carry out a detailed investigation of major functional properties, including bacterial killing, by PMN from patients with severe sepsis or septic shock and to relate those observations to the clinical status of the patients. PMN showed increased basal O_2_^.-^ production, decreased phagocytosis-related ROS production, and impaired elimination of *Staphylococcus aureus* that correlated with sepsis severity. Peptides in the range of 3-12 kDa isolated from the blood of septic patients induce the same functional changes in healthy neutrophils. Both the total concentration of peptides in the blood plasma of septic patients and the functional impairment caused by the peptides are proportional to the clinical status of the patients. Reduction of the peptide concentration in septic patients could provide a potential target for intervention.

## Results

### Altered effector functions of PMN isolated from septic patients

To determine the relationship between severity of sepsis and neutrophil dysfunction, unstimulated, maximal and bacteria-evoked ROS production, phagocytosis and antibacterial capacity were measured in neutrophils from septic patients and compared to Acute Physiology and Chronic Health Evaluation (APACHE II) score at the time of blood sampling (Fig.1). Unstimulated extracellular superoxide (O_2_^.-^) release was five times higher in PMN from septic patients than from healthy donors, and the individual values showed significant positive correlation with the APACHE II score, i.e. worsening of the clinical status of the patient (Fig.1a). Maximal O_2_^.-^ production elicited by the pharmacological agent phorbol-myristate-acetate (PMA) was significantly lower in septic than in healthy PMN, but no correlation was found with the clinical score (Fig.1b). ROS production elicited by phagocytosis of Gram positive bacteria *S. aureus* was significantly lower in PMN from septic patients than from healthy donors and showed a negative correlation with the APACHE II score (Fig.1c). Similar data were obtained with phagocytosis of Gram negative bacteria, *E. coli* (Fig.1d). Phagocytic capacity was tested at two different loads of bacteria. We previously reported that the higher load approaches the maximal capacity of neutrophils under our conditions (26). No difference in phagocytosis was detected between neutrophils from healthy or septic donors at either load (Fig.1e). Survival of *S. aureus* was significantly increased (i.e. bacteria elimination was decreased) when a higher load of bacteria were incubated with septic PMN, and the values showed significant positive correlation with the APACHE II score (Fig.1f). No significant differences in bacterial survival and no correlation with the clinical status were observed following incubation with a lower dose of *S. aureus* or with either dose of *E. coli* (Fig. 1g-i), although scattering of the data was larger than in PMN from healthy subjects. Thus, alterations in basal ROS release, phagosomal ROS generation, and killing of *S. aureus* under conditions of high bacterial load correlated with the severity of sepsis.

**Fig. 1.:**
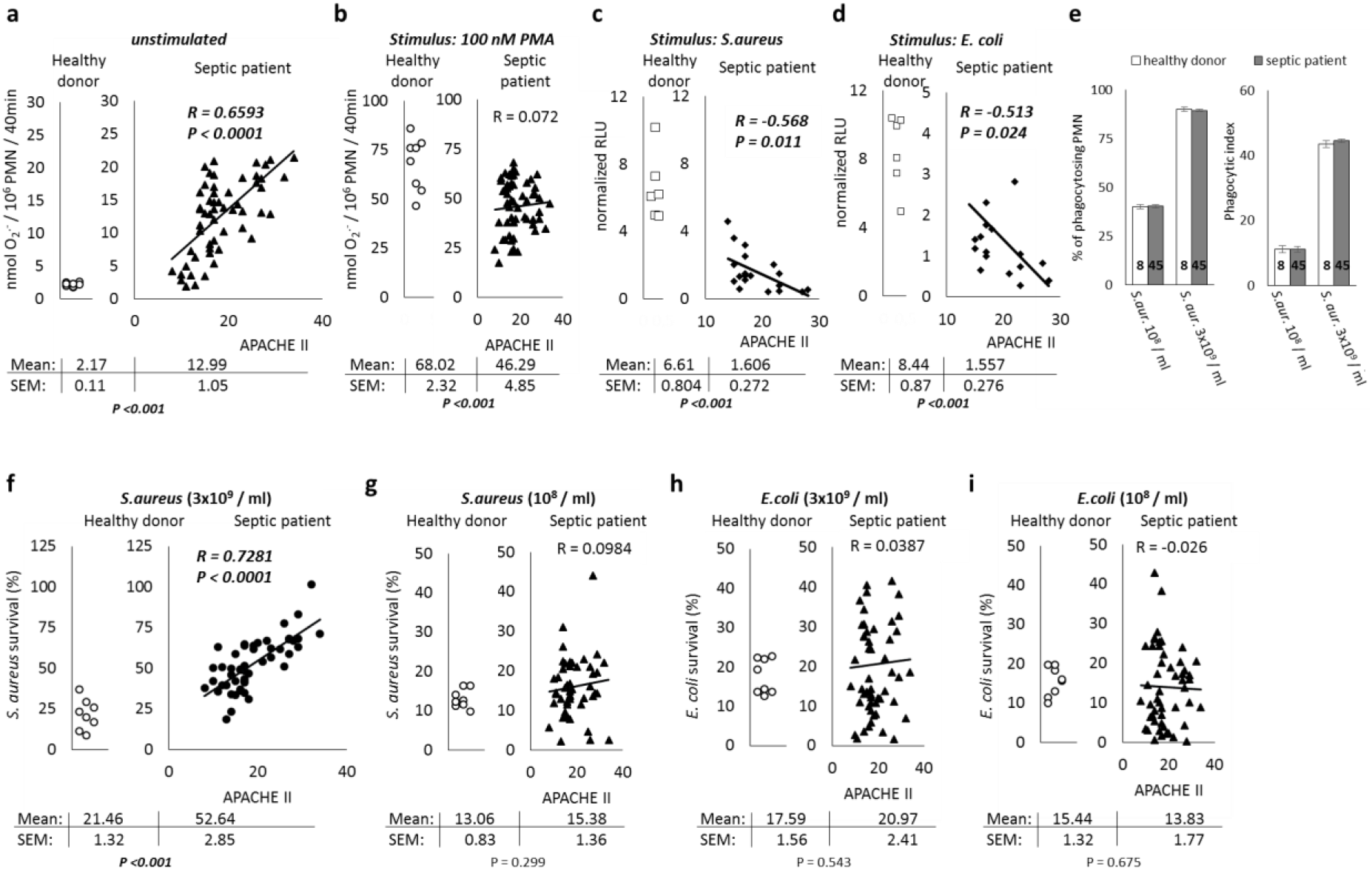
Functional properties of PMN from septic patients. Unstimulated (a.) or 100 nM PMA stimulated (b.) extracellular superoxide release by PMN from healthy and septic donors, measured by SOD-inhibitable cytochrome c reduction assay, correlated to APACHE II score. c., d.: lucigenin-based intracellular ROS production during phagocytosis of *S. aureus* (c.) or *E. coli* respectively (d.), correlated to APACHE II score. e.: ratio of phagocytosing PMN to all PMN and phagocytic index of PMN from healthy and septic donors respectively with the marked concentrations of *S. aureus*. f., g.: Survival of marked amount of opsonized *S. aureus* after incubation with PMN from healthy or septic donors respectively, related to APACHE II score. h., i.: Survival of marked amount of opsonized *E. coli* after incubation with PMN from healthy or septic donors respectively, related to APACHE II score. R and P numbers in *italic style* refer to significant correlation. Tables under curves show mean and SEM with level of significance. In panel e. error bars represent SEM, indicated numbers in columns represent number of independent samples. Significance above p > 0.05 is not indicated.

To determine the relationship of altered PMN responses to clinical outcome, unstimulated O_2_^.-^ release and survival of larger doses of *S. aureus* were compared among specific groups of patients. Both parameters differed significantly between patients who survived or died during the observation period (Suppl.Fig.1a), with survivors showing reduced basal O_2_^-^ release and enhanced ability to kill bacteria. In contrast, no differences were found in either parameter in patients with Gram negative, Gram positive, or no bacteria identified in blood cultures (Suppl.Fig.1b). Similarly, no significant differences were detected based on the location of the infection (Suppl.Fig.1c).

Next, we tested whether alterations of the extracellular environment typically occurring in septic states, such as acidic or alkaline pH, hyperthermia, hyperglycaemia or a combination of these, may cause any of the observed changes of neutrophil functions (Suppl.Fig.2). No significant changes were detected in unstimulated, maximal, or bacteria-induced ROS production or survival of either of the two test bacteria. Thus, neutrophil dysfunction in septic patients is related to the disease, not to changes in the internal environment typically occurring in sepsis.

### Effect of septic plasma on PMN from healthy donors

To determine if soluble factor(s) in blood plasma altered neutrophil functions, the effect of incubating PMN from healthy donors with blood plasma from septic patients was examined. Neither pathogens nor antibiotics were present in the stored septic plasma (data not shown). PMN from healthy donors were incubated for one hour in plasma (two times diluted with PBS) from septic patients or healthy individuals or in the absence of plasma, and PMN functions examined after washing the plasma away and resuspending PMN in HBSS. Based on binding of annexin V and propidium-iodide, septic plasma had no effect on the viability of healthy neutrophils (data not shown).

Extracellular O_2_^.-^ release by unstimulated healthy PMN was significantly increased by pretreatment with septic plasma, while maximal release of O_2_^-^ stimulated by PMA was significantly decreased (Fig.2a). There was a significant positive correlation between O_2_^.-^ release by septic plasma-treated healthy PMN and the clinical status of the patient (Fig.2b). On the other hand, the effect of plasma from septic patients on maximal O_2_^.-^-production of healthy PMA-activated PMN was not related to the clinical status (Fig.2c).

**Fig. 2.:**
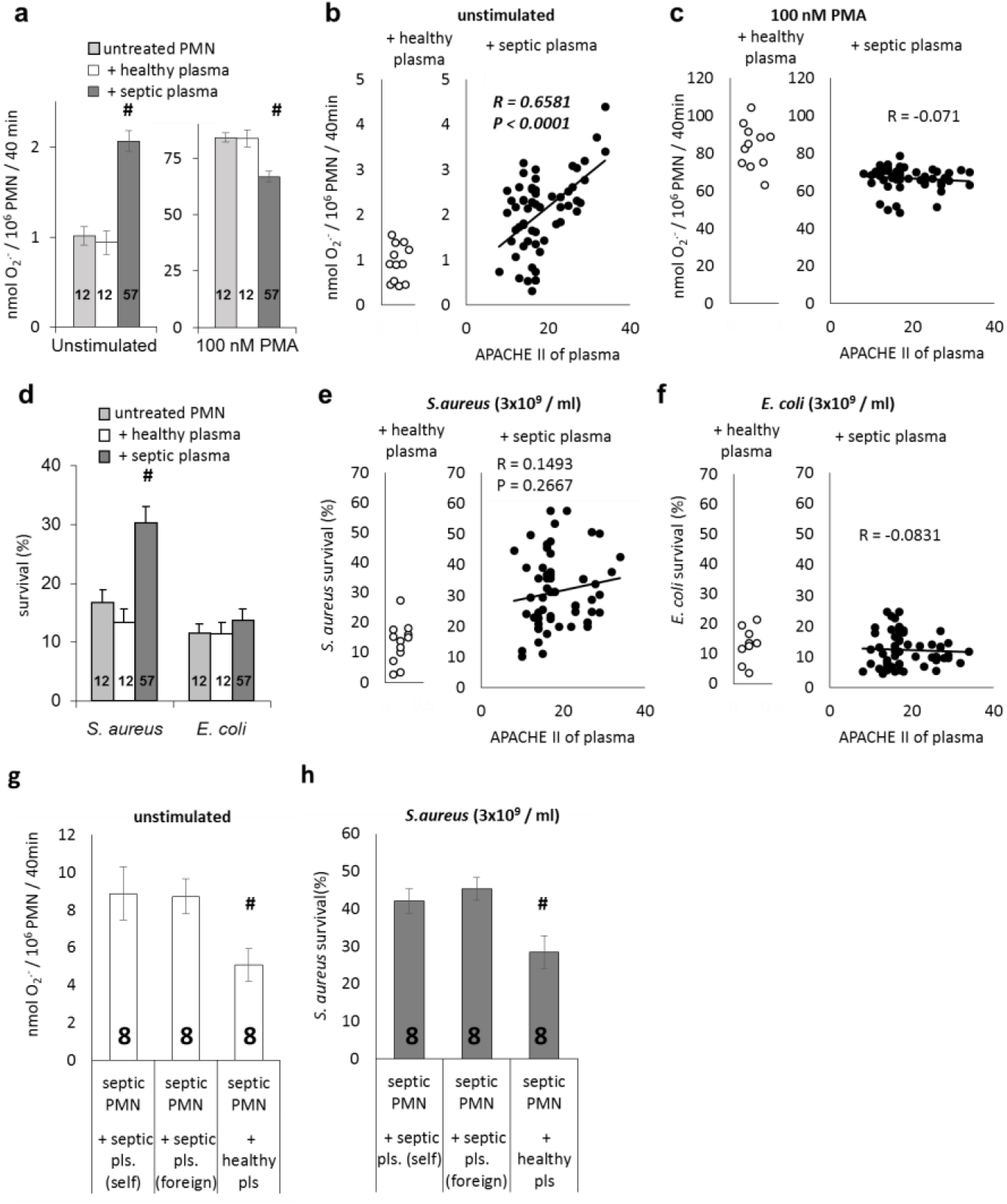
Functional properties of PMN from healthy donors treated with blood plasma from septic patients. a.: Unstimulated or 100 nM PMA stimulated extracellular superoxide release by PMN from healthy donors, either without treatment or following treatment with blood plasma from healthy or septic donors. b., c.: correlation of extracellular superoxide release to APACHE II score of the plasma donor. d.: survival of 3×10^9^ / ml *S. aureus* or *E. coli* after incubation with PMN from healthy donors, either without treatment or following treatment with blood plasma from healthy or septic donors. e., f.: correlation of bacterial survival to APACHE II score of plasma donor patients. g., h.: Effect of blood plasma from septic patients (self or foreign respectively) or from healthy donors on unstimulated superoxide release (g.) and on antibacterial effect (h.) of PMN from septic patients. Error bars represent SEM. Indicated numbers in columns represent number of independent samples. #: p < 0.01. R and P numbers in *italic style* refer to significant correlation. Significance above p > 0.05 is not indicated.

Incubation of healthy PMN with plasma from septic patients significantly decreased *S. aureus* killing (Fig.2d and e), while there was no difference in survival of *E. coli*. (Fig.2f). The phagocytic activity of PMN incubated with blood plasma from septic patients or healthy donors was not significantly different (data not shown).

To determine reversibility of the functional alterations, PMN isolated from septic patients were incubated for one hour in autologous plasma, in plasma from another septic patient, or in plasma from a healthy donor, then washed and resuspended in PBS. Figs.2g and 2h show that unstimulated O_2_^.-^ release and survival of *S. aureus* were significantly reduced upon incubation of septic PMN in healthy plasma. Taken together, our data show that plasma from septic patients reproduced the same pattern of functional changes in healthy PMN as observed in PMN from septic patients. The altered activity in PMN from septic patients was reversed upon replacement of septic plasma by normal plasma. These results suggest that PMN dysfunction in sepsis is induced by soluble plasma factor(s).

### Characterization of the plasma factor(s)

To characterize the plasma factor(s) that induce altered function in septic PMN, we pre-treated septic plasma with dialysis, heat and acid inactivation, and various enzymes prior to incubation with healthy PMN. As shown in Figs.3a-c, incubation of healthy PMN with plasma from septic patients again induced enhanced survival of *S. aureus*, increased unstimulated extracellular O_2_^.-^ release, and reduced maximally stimulated O_2_^.-^ release. Dialysate from septic plasma containing peptides < 12 kDa reproduced the changes in *S. aureus* survival and O_2_^.-^ release seen in septic PMN and in healthy PMN incubated with septic plasma. On the other hand, incubation of healthy PMN with septic plasma from which peptides <12 kDa were removed by dialysis did not reproduce any of those changes in PMN function. Heat treatment at 100°C for 15 min or treatment with 0.1 M acetic acid for 20 min did not affect the ability of septic plasma to alter functional responses of healthy PMN (Figs.3d-f).

**Fig. 3.:**
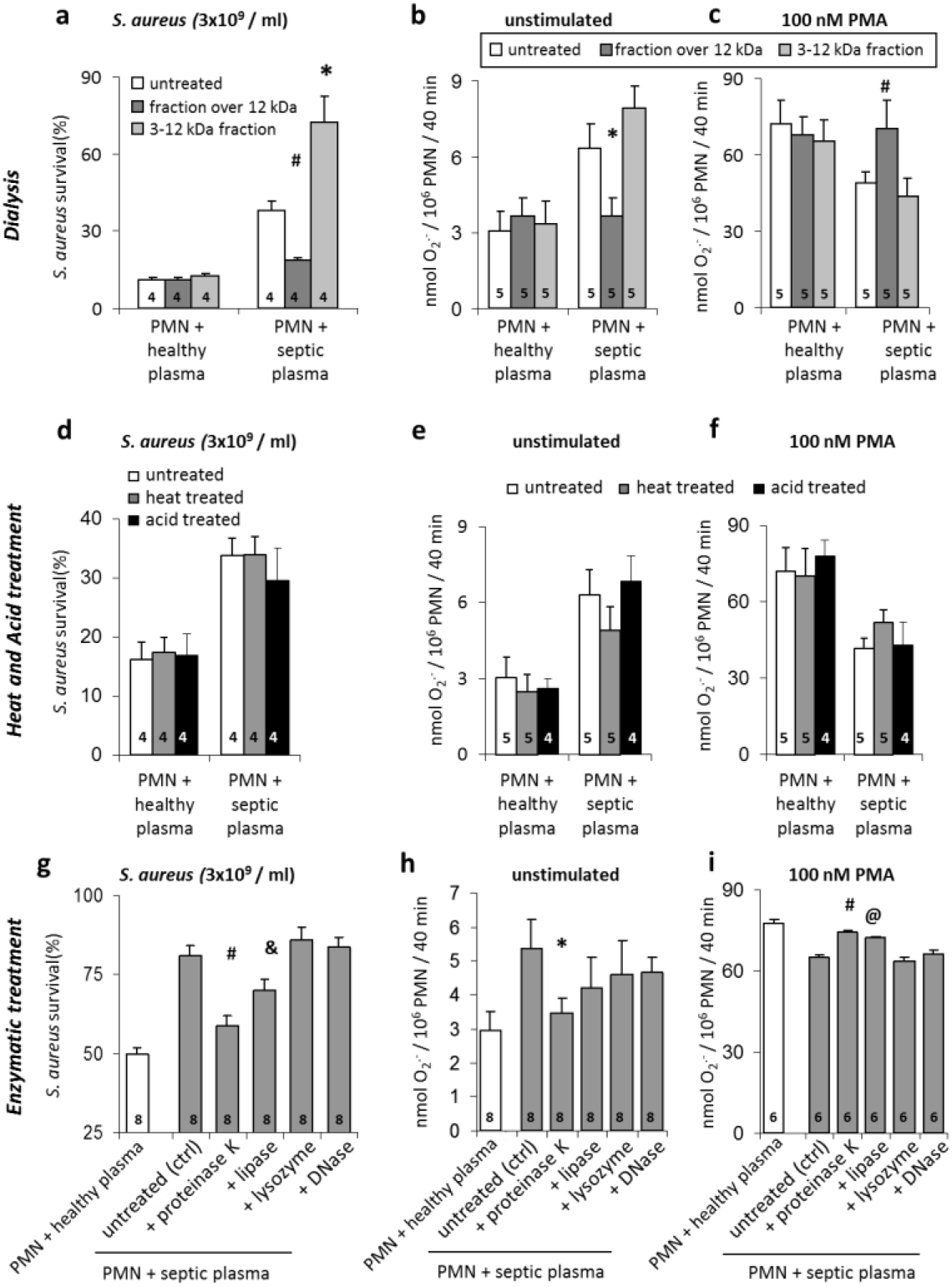
Effect of treated fractions of blood plasma from healthy or septic donors on healthy PMN. a. - c.: Effect of dialyzed blood plasma fractions on (a.) *S. aureus* survival (3×10^9^ CFU/ml) and on superoxide release by (b.) unstimulated or (c.) PMA stimulated PMN from healthy donors. d.-f.: Effect of heat or acid treated blood plasma on (d.) *S. aureus* survival (3×10^9^CFU/ml) and on superoxide release by (e.) unstimulated or (f.) PMA stimulated PMN from healthy donors. g.-i.: Effect of enzymatically treated blood plasma on (g.) *S. aureus* elimination (3×10^9^CFU/ml) and on superoxide production of (h.) unstimulated or (i.) PMA stimulated PMN from healthy donors. Error bars represent SEM. Indicated numbers in columns represent number of independent samples. #: p < 0.01, *: p < 0.05, &: p = 0.26, @: p = 0.101 compared to untreated plasma samples. Significance above p > 0.05 is not indicated.

Next, we tested the effect of selected enzymes on the septic plasma (Figs.3g-i). Treatment with proteinase K resulted in almost complete loss of the effect of septic plasma on healthy PMN. Porcine pancreatic lipase (which may contain protease as impurity) had a small, but non-significant, effect. Lysozyme and DNase had no effect on the ability of septic plasma to alter functions of healthy PMN. Similar data were obtained when the concentrated dialysate of septic plasma containing peptides <12 kDa was subjected to enzyme treatment (data not shown). Our data suggest that septic plasma contains heat- and acid-resistant peptide(s) with a molecular size between 3 and 12 kDa that induce altered PMN activity *in vitro* and *in vivo*.

### Relation between the amount of dialyzable peptides and impairment of neutrophil functions

To determine whether the amount of dialyzable peptides in the plasma of septic patients is related to PMN dysfunction, the quantity of dialyzable peptides in selected plasma samples from septic patients was compared to clinical and functional data shown in Figs.1a and 1f. Fig.4a compares unstimulated PMN extracellular O_2_^.-^ release to the amount of dialyzable peptides in each patients’ plasma sample. Fig.4b shows a similar analysis for *S. aureus* survival in the presence of PMN from septic patients. Both PMN parameters showed a statistically significant positive correlation with the peptide concentration in plasma from septic patients. Next, the amount of dialyzable peptides was compared to the APACHE II score for each patient (Fig.4c). This correlation proved to be highly significant. Thus, circulating peptides are potential candidates for impairing neutrophil functions in septic patients.

**Fig. 4.:**
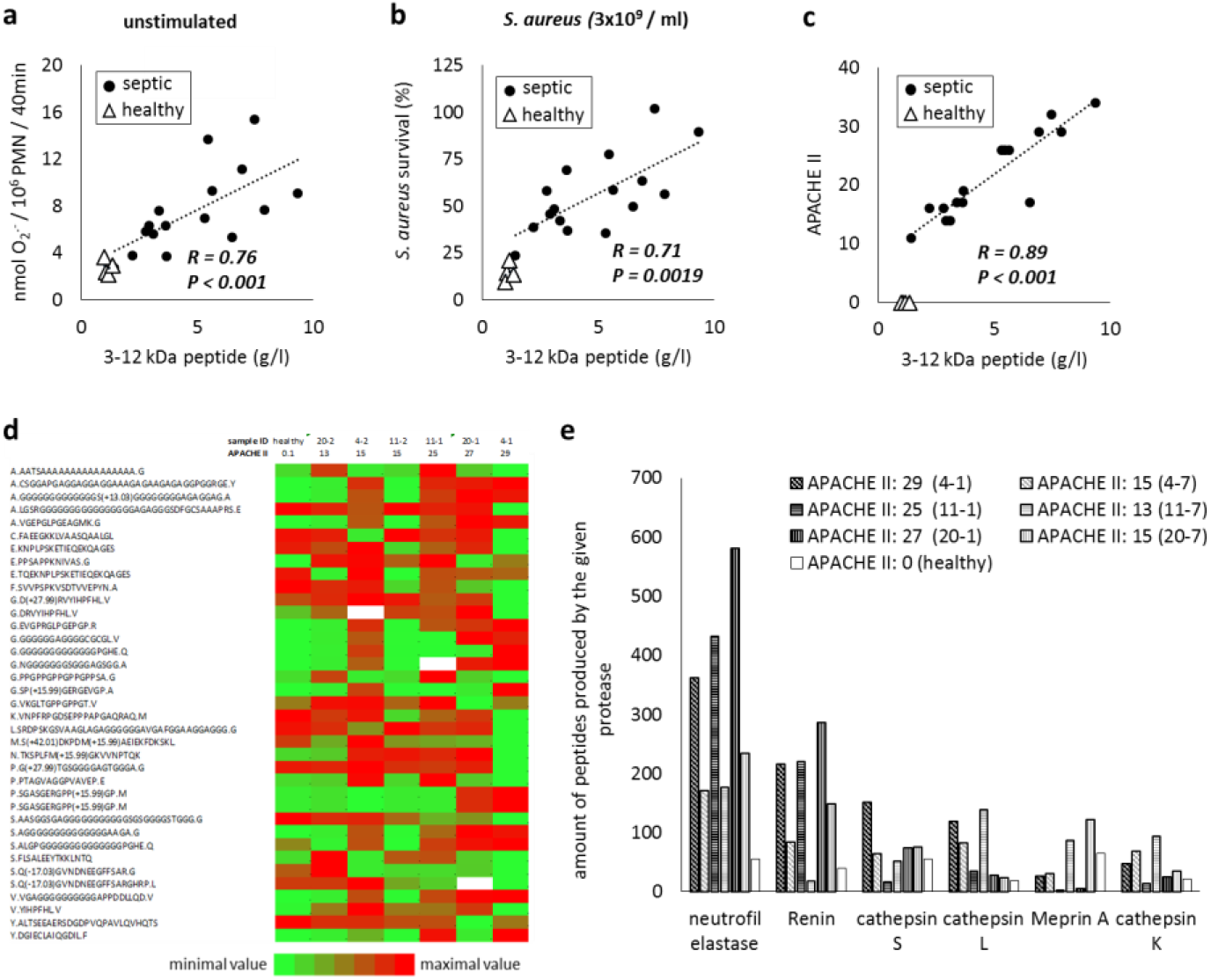
Correlation of peptide content of the 3-12 kDa fraction of blood plasma with functional parameters of PMN from and clinical status of septic patients. Correlation of peptide content of 3-12 kDa fraction of blood plasma with a.: unstimulated superoxide release, and with b.: antibacterial (3×10^9^CFU/ml *S. aureus*) effect of PMN from septic patients. c.: with APACHE II score of the patient. R and P numbers in *italic+bold style* refer to significant correlation. d.: Amount of 30 most common peptides, found by proteomics analysis in 3-12 kDa fraction of two samples of each of three patients and of a healthy sample. Color code ranges from smallest to highest value of given peptide across the seven samples. White color means no detection in the given sample. Since values vary in two orders of magnitude, logarithmic scale was used. e.: Five most common proteases involved in peptide production, based on the amount of peptides generated by them in the seven samples mentioned above. Numbers in parenthesis show patient ID, the first number indicating the patient, the second the time of sampling (earlier/later).

### Proteomic analysis of the peptide composition of septic plasma

To identify the peptide(s) responsible for impairment of neutrophil functions, proteomic analysis of selected plasma samples was performed. We chose 3 pairs of samples drawn from patients on the 1^st^ and 7^th^ day of observation. The APACHE II scores on the 1^st^ and 7^th^ day were 29/15, 25/13 and 27/15. One sample from a healthy volunteer was analyzed in parallel. Peptides with molecular sizes between 3 and 12 kDa were identified from plasma dialysate and quantified by label-free analysis of peptide peaks. A total of 214 peptides from 88 unique proteins were present in at least 6 out of the 7 samples (Suppl.Table1). Those proteins consisted of plasma proteins and proteins primarily localized in cells or the extracellular matrix. The proportion of peptides derived from proteins in those different locations varied randomly among the patients and by clinical status. One common phenomenon was a higher number of peptides derived from cellular proteins in the samples taken on the 1^st^ day of observation, when the APACHE II score of the patient was higher (data not shown). No fragments of cytokines or chemokines were identified. Using OneDCRF (Uni. Umea, Sweden) web-based program and ModiDB-lite program (University of Padova, Italy) for the analysis of peptide structure, peptides showed a high frequency (72 to 98%) of disordered structure, consistent with the heat and acid resistant properties of the dialyzed peptides (Fig.3d to f) (27–30).

A quantitative comparison of the 30 most abundant peptides in patient dialysates is shown by the heat map in Fig. 4d. None of the peptides showed a consistent quantitative change with increasing APACHE II score. Similar results were found with analysis of the less abundant peptides (raw data in Suppl.Table1). The lack of correlation of the amount of any individual peptide with APACHE II score contrasts to the linear relationship observed between the total amount of peptides and the APACHE II score shown in Fig.4c (which also was evident for the 7 samples investigated by proteomic analysis).

To identify potential enzymes generating peptides in septic plasma, protease prediction with Proteasix (www.proteasix.org) was performed on each peptide, using only experimentally observed cleavage sites. The proteases predicted to generate the most peptides are presented in Fig.4e, and similar analysis containing predicted proteases for the remainder of less common peptides is shown in Suppl.Fig.3a. The largest number of identified peptides were produced by neutrophil elastase (NE) and renin. Although the amount of peptides with the relevant signature sequence showed great difference in the individual samples, in all 3 patients the amount of peptides was higher in the 1^st^ day samples than in samples taken on the 7^th^ day and exceeded the amount of the healthy sample. The amount of less abundant peptides carrying the target sequence of other proteases also showed large variations between patients and clinical status (Suppl.Fig.3a).

### Possible origin of the transfer factor(s)

Analysis of peptides identified in septic plasma indicated both cellular and plasma proteins could serve as a source of peptides affecting neutrophil functions. This observation lead us to search different cell lysates for the ability to alter PMN function. PMN of healthy donors were incubated with dialysate (3-12 kD) and non-dialyzable (above 12 kD) fractions of bacteria and various blood cells and plasma (Suppl.Fig.3b and c). The pattern of effects produced by *S. aureu*s lysates differed from that observed with plasma from septic patients, as the non-dialyzable fraction brought about a definitive impairment of neutrophil functions, whereas the dialysate had a weaker effect. In contrast, the dialysate from human PMN lysates increased both unstimulated extracellular O_2_^.-^ release by PMN from healthy donors and survival of *S. aureus*, indicating impaired elimination capacity. Fractions of lysates of monocytes, lymphocytes and red blood cells as well as plasma of healthy patients had no effect on PMN functional responses. Thus, cells and plasma do not normally contain peptides capable of altering PMN function.

Next, we tested the possibility that proteases contained in PMN lysates produced dialyzable peptides that impair PMN functions. Lysates of peripheral blood mononuclear cells (PBMC) that did not alter PMN functions were treated with proteinase K or PMN lysate, and the formation of dialyzable peptides (Fig.5a) and their effect on neutrophil functions (Fig.5b and c) followed over time. The representative experiments shown indicate that the time of maximal increase of unstimulated O_2_^.-^ release (Fig.5b) and bacterial survival (Fig.5c) coincides with the time of maximal production of dialyzable peptides for both proteinase K and PMN lysate (Fig.5a). Figs.5d and e summarize data showing that unstimulated O_2_^.-^ release (Fig.5d) and *S. aureus* survival (Fig.5e) are significantly correlated with the amount of dialyzable peptides generated by both protease treatments. Similar results were obtained with lysates of red blood cells and plasma as substrates (data not shown).

**Fig. 5.:**
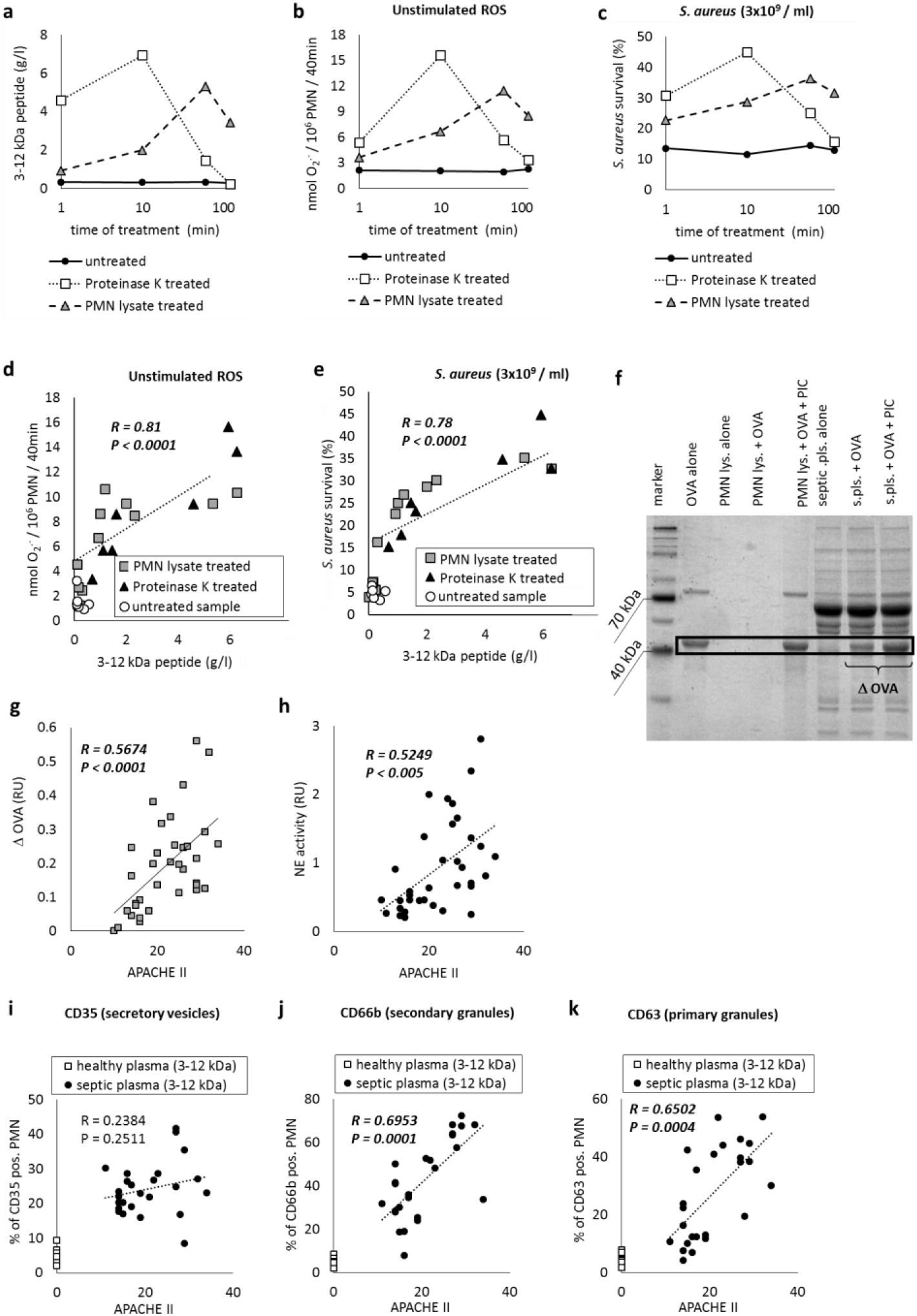
Origin of the 3-12 kDa peptide fraction of blood plasma from septic patients. a.-c.: Representative sample of enzymatic digestion of a PBMC lysate. a.: Effect of different enzymes on peptide content of 3-12 kDa fraction of PBMC in time. b.: Effect of 3-12 kDa fraction collected at the given time point on unstimulated superoxide release and c.: on antibacterial properties (against 3×10^9^CFU/ml, opsonized *S. aureus*) of naïve PMN from healthy donor. d., e.: Correlation of peptide content of 3-12 kDa fraction of enzymatically digested lysates of PBMC or RBC or blood plasma with effect of these fractions on d.: unstimulated superoxide release and on e.: antibacterial properties (3×10^9^CFU/ml, opsonized *S. aureus*) of PMN from healthy donors. Correlation was calculated on the basis of all data obtained from digestions with different enzymes and lysates. f.-h.: Proteolytic activity of plasma of septic patients. f.: representative SDS-PAGE gel electrophoresis of different samples after incubation with ovalbumin (OVA). PIC = protease inhibitor cocktail. s.pls. = blood plasma from septic patient. ΔOVA= difference in Coomasie density of OVA in the presence and absence of PIC. g.: correlation of APACHE II score with ΔOVA. h.: correlation of APACHE II score with neutrophil elastase (NE) activity. i.-k.: Effect of 3-12 kDa fraction of blood plasma on degranulation of naïve PMN. Correlation of APACHE II score with ratio of i.: CD35 positive (secretory vesicles), j.: CD66b positive (secondary granules), and k.: CD63 positive (primary granules) naïve PMN after incubation with 3-12kDa fraction of blood plasma of septic patient. R and P numbers in *italic style* refer to significant correlation. Significance above p > 0.05 is not indicated.

Next, we addressed whether the clinical status of the patients was related to the protease activity in plasma by two different techniques. First, the total protease activity in plasma was assessed by determining protease inhibitor sensitive reduction in ovalbumin (OVA) digestion by SDS-PAGE. A representative experiment is shown in Fig.5f. The summarized data, presented as the difference in the intensity of the OVA band in the presence and absence of protease inhibitor cocktail, show a significant positive correlation with the clinical score (Fig.5g). Second, we determined NE activity in the plasma of the septic patients (Fig.5h). NE activity showed a significant positive correlation with the APACHE II score.

To determine if PMN exocytosis was the source of plasma NE in septic patients, the extent of exocytosis of different granule subsets induced by incubation of healthy PMN with plasma dialysate from healthy and septic patients was determined (Fig.5i to k). The percent of cells with enhanced plasma membrane expression of CD66b (a marker for secondary granules) and CD63 (a marker of primary granules) was significantly correlated with the APACHE II score of patients from whom plasma was obtained (Fig.5j and k). Plasma dialysate from healthy subjects failed to induce increased expression of either marker. Although a higher percent of PMN incubated with septic plasma dialysate expressed CD35 (a marker of secretory vesicles), no significant correlation was found between the APACHE II score and CD35 expression (Fig.5i). This may be explained by the ability of secretory vesicles to undergo complete exocytosis at low concentrations of stimuli(31–33). Taken together, our data suggest that protease release from PMN is stimulated by peptide(s) in the 3-12 kDa size range of septic plasma, and those proteases generate peptides that alter the effector functions of PMN.

### Role of intracellular ROS generation in alteration of PMN functions in septic patients

Our data show that PMN from septic patients release increased O_2_^.-^ into the extracellular space, but show decreased intracellular ROS production upon bacterial phagocytosis (Fig.1). This discrepancy initiated a more detailed investigation of localization of ROS production using C400, an intracellularly trapped fluorescent ROS detector, to quantify the effect of peptides on intracellular ROS generated in response to phagocytosis. Healthy PMN exhibited a high level of intracellular ROS production when treated with 3-12 kDa fraction of healthy plasma prior to phagocytosis of opsonized *S. aureus* for 20 min (Fig.6a and Suppl.Video1). Pretreatment of the same cells with 3-12 kDa fractions of septic plasma (Fig.6b) or with 3-12 kDa fractions of protease-treated cell lysate (Fig.6c and Suppl.Video2) resulted in an obvious reduction in intracellular ROS production during phagocytosis.

**Fig. 6.:**
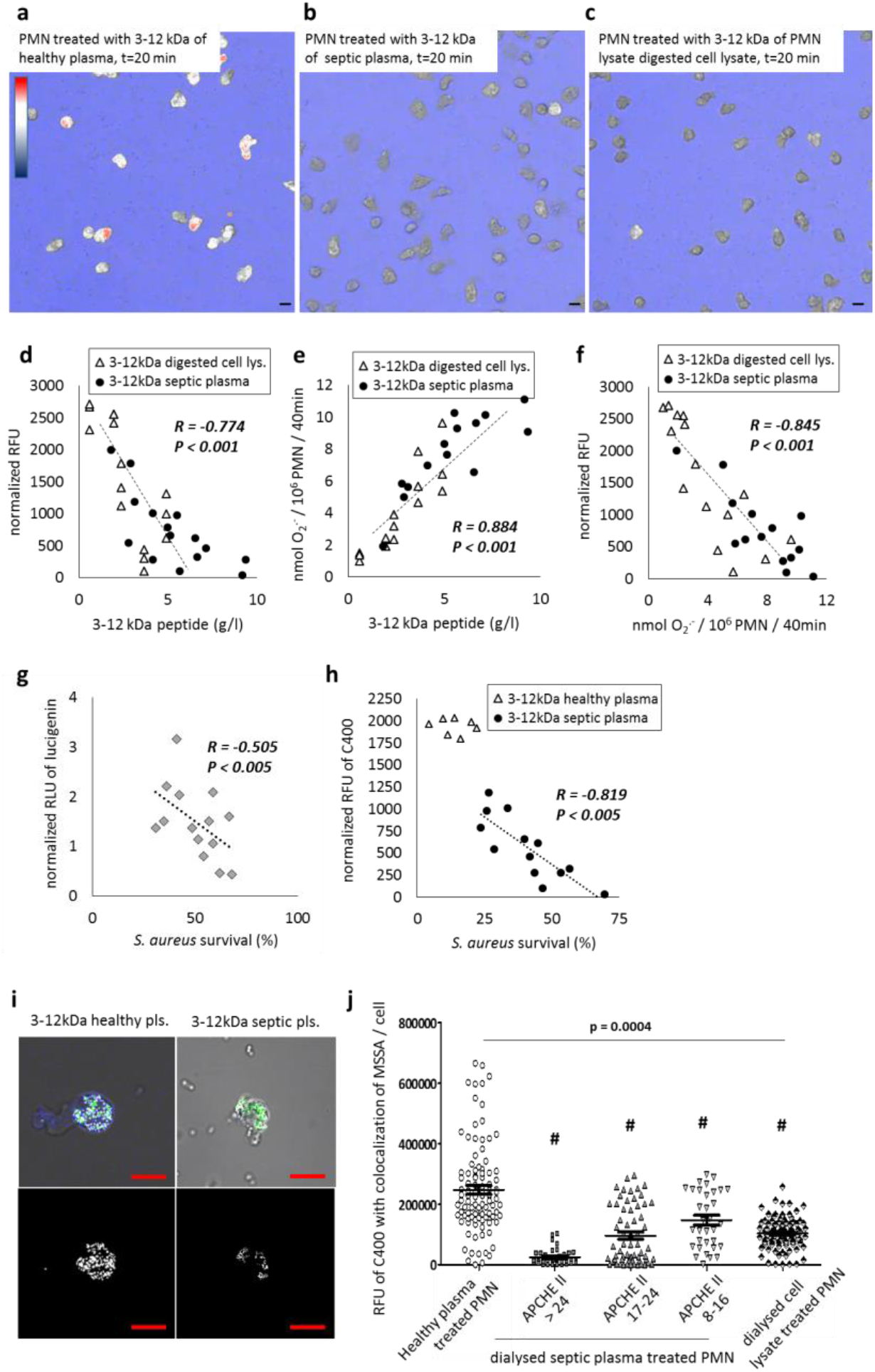
Analysis of intracellular ROS production of PMN. a.-c.: Representative pictures of C400 fluorescence indicating intracellular ROS production of PMN from healthy donors treated previously with 3-12 kDa fraction of a.: plasma from healthy donors, b.: plasma from septic patients, c.: PMN lysate-digested PBMC lysate, after 20 min incubation with opsonized *S. aureus*. d.: correlation of intracellular ROS production of PMN from healthy donors, treated with 3-12 kDa fraction of plasma from septic patients or of digested cell lysates, after 20 min incubation with opsonized *S. aureus* to the peptide content of the used fraction. e.: correlation of unstimulated extracellular superoxide production of PMN from healthy donors, treated with the 3-12 kDa fraction of plasma from septic patients or of digested cell lysates to the peptide content of the used fraction. f.: correlation of intracellular ROS production during phagocytosis (data from panel d.) to unstimulated extracellular superoxide release (data from panel e.). g., h.: Correlation of antibacterial properties (against 3×10^9^CFU/ml *S. aureus*) to phagocytosis-induced intracellular ROS productions of PMN from g.: septic patients (measured by the lucigenin assay) or h.: PMN from healthy donors treated with 3-12 kDa fractions of plasma from healthy or septic donors (measured by C400 assay). i., j.: ROS production around phagocytosed bacteria. i: Representative confocal microscopic picture of phagocytosing PMN from healthy donors, pretreated with 3-12 kDa fraction of plasma from healthy or from septic donors. Blue: C400. Green: GFP expressing opsonized *S. aureus* (USA300). White: colocalization of C400 and bacteria. Upper row: merge of all channels with phase contrast image. Lower row: colocalization layer of the same image. j.: Relative fluorescence intensity (RFU) of C400 at the sites of colocalization with bacteria in a total of 100 PMN/category, of 4 independent experiments. Error bars represent SEM. #: p < 0.01. R and P numbers in *italic style* refer to significant correlation.

Next, intracellular ROS production was compared to the concentration of dialyzable peptides with which phagocytosing PMN were pretreated. Significant negative correlations were found for the 3-12 kDa fraction of plasma from septic patients and for the 3-12 kDa fraction from protease-digested PBMC lysate (Fig.6d). Values from both fractions fall on the same line. Basal extracellular O_2_^.-^ release showed a significant positive correlation with peptide concentration using combined data points obtained with fractions of septic plasma and of digested PMN lysate (Fig.6e). Additionally, intracellular ROS production showed a significant negative correlation with extracellular O_2_^.-^ release (Fig.6f).

Elimination of *S. aureus* is strictly dependent on ROS production (34). Therefore, we compared phagocytosis-induced ROS production to *S. aureus* survival in PMN from septic patients (Fig.6g) and in PMN from healthy subjects preincubated with 3-12 kDa fractions of septic or healthy plasma (Fig.6h). In both cases we obtained a significant negative correlation indicating that a lower intracellular ROS production was associated with a higher *S. aureus* survival. To compare localization of phagocytosed bacteria and intracellular ROS production, confocal microscopic images of C400-loaded PMN after phagocytosis of green fluorescent protein expressing bacteria were obtained. Fig. 6i shows colocalization of the two dyes, as indicated by the white dots on merged images. Difference in C400 fluorescence co-localization with phagocytosed bacteria between PMN treated with the 3-12 kDa fraction from blood plasma of healthy donors (left panels) or septic patients (right panels) was visually obvious. For statistical analysis, the average intensity of C400 fluorescence colocalization with green fluorescent bacteria was determined (Fig.6j). That intensity value was significantly higher in PMN treated with the dialyzable plasma fraction from healthy donors, compared to PMN treated with 3-12 kDa fraction of plasma from septic donors or 3-12 kDa fraction of digested cell lysates. Additionally, intracellular ROS production colocalized with bacteria decreased as the APACHE II score of patients from whom plasma samples were obtained increased. Thus, increased extracellular O_2_^.-^ release by PMN from septic patients is associated with decreased ROS production within phagosomes and decreased bacterial killing.

### Cellular damage brought about by septic plasma treated PMN

Based on enhanced granule exocytosis and increased extracellular release of ROS by PMN from septic patients, we postulated that PMN exposed to plasma from septic patients damage bystander cells. To test this hypothesis, various intact blood cells were incubated with PMN pretreated with 3-12 kDa fractions of plasma from healthy donors or septic patients or with dialysates obtained from digested cell lysates (Fig.7). Cell damage was measured by staining with annexin V and propidiumiodide (PI), two characteristic markers of upcoming cell death. The proportion of intact (not stainable with either annexin V or PI) cells (PBMC or PMN) decreased to 50 to 60% upon incubation with PMN pretreated with either 3-12 kDa fractions of septic plasma or digested dialyzed cell lysate (Fig.7a,b). Similar results were obtained when Erytrosin B positivity of erythrocytes (RBC) was investigated (Fig.7c). In a complementary approach, release of cytosolic proteins was measured. A 3 to 5-fold increase in lactate dehydrogenase (LDH) release from PBMC or PMN was detected after incubation with PMN pretreated with 3-12 kDa fraction of septic plasma or digested, dialyzed cell extract (Fig.7a,b). A significant release of hemoglobin from erythrocytes occurred under identical conditions (Fig.7c). We interpret these data to indicate that PMN activated by peptides in the plasma of septic patients damage bystander cells.

**Fig. 7.:**
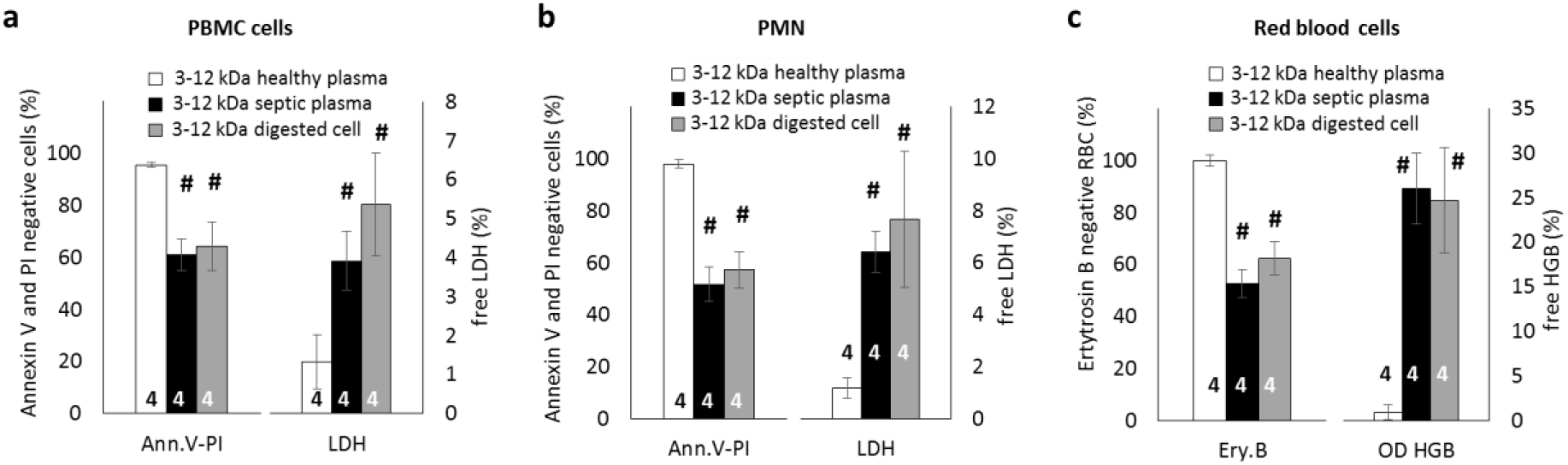
Effect of activated PMN on other blood cells. Effect of 10^6^ / ml PMN from healthy donors, pretreated with 3-12 kDa fractions of plasma from healthy or septic donors or PMN lysate-digested cell lysate respectively, on viability and integrity of a.: 10^7^ / ml PBMC, of b.: 10^7^ / ml PMN and of c.: 10^9^ / ml RBC from the same donor. Viability and integrity were characterized in case of PBMC and PMN by Annexin V and propidium iodide negativity (left) and LDH release (right) and for RBC by erytrosin B positivity (left) and hemoglobin (HGB) release (right). (n = 4, ±SEM, #: p < 0.01 compared to PMN treated with 3-12 kDa fraction of plasma of healthy donor. Significance above p > 0.05 is not indicated.

## Discussion

The major finding of the present study is that peptides in the blood of septic patients induce neutrophil dysregulation, leading to impaired bactericidal activity and enhanced injury to normal cells. Our data show altered functional responses in PMN isolated from septic patients, including enhanced basal extracellular release of ROS, impaired intra-phagosomal ROS generation, and reduced bactericidal activity. Incubation of normal PMN with plasma from septic patients or peptides in the molecular size range of 3-12 kDa contained in that plasma reproduced the abnormal functions observed in septic PMN. The significant relationship of altered PMN functional responses, peptide concentration in the blood, and proteolytic activity in the blood to the severity of illness, determined by the APACHE II score, supports the clinical relevance of our findings.

Several lines of evidence suggest that a pool of different peptides, rather than one specific peptide, are responsible for the alterations in PMN function. First, the concentration of all peptides showed a significant correlation with the APACHE II score (Fig.4c), while the proteomic analysis failed to identify any peptides showing a significant correlation (Fig.4d). Second, neutrophil-activating peptides were produced from very different substrates, including cell lysates, plasma proteins and bacterial lysates. Third, different enzymes were able to produce peptides exerting the same effects on neutrophil function (Fig.5d and e). Fourth, proteomic analysis identified peptides generated by several enzymes in the plasma of septic patients, and those peptides were present in different proportions among individual patients and when categorized by clinical status (Fig.4e and Suppl.Fig.3a).

Our data indicate that PMN serve as both the target of circulating peptides in sepsis, and as the source of at least some of the proteases generating those peptides. Our results showed that secretory vesicles, specific granules, and azurophilic granules undergo mobilization to the plasma membrane upon exposure to plasma dialysate from septic patients. Thus, peptides generated from proteins in quite different locations and by different proteases stimulate PMN extracellular release of granule cargo containing numerous proteolytic enzymes likely involved in further peptide generation. Indeed, the activity of neutrophil elastase in septic samples was elevated in proportion to severity of the disease.

The mechanism by which circulating peptides generated in response to sepsis induce disparate PMN functions remains to be determined. Disordered peptides are characterized by high flexibility and accessibility, suggesting many potential reaction partners (35). The observed changes in neutrophil granule exocytosis provide one possible explanation for enhanced extracellular ROS release and bystander cell injury and reduced phagosomal ROS production and impaired bactericidal activity. Granule mobilization leading to translocation of membrane components of the NADPH oxidase to the phagosomal and plasma membranes is necessary for enhanced ROS generation associated with PMN priming (36). Generation of ROS and release of toxic granule cargoes act synergistically to kill bacteria within phagosomes and to injury cells when released extracellularly (37–39). These observations raise the possibility that circulating peptides misdirect PMN granules to fuse with the plasma membrane. This event would result in enhanced release of both ROS and granule cargo, increasing the risk of injury to bystander cells. Direct peptide inhibition of granule fusion with phagosomes or limiting the number of granules available to phagosomes through enhanced targeting to plasma membranes would result in decreased phagosomal content of ROS and anti-microbial granule cargo, leading to impaired anti-microbial activity. Thus, we interpret our results to indicate that neutrophils are the central component of a feedback loop in sepsis that impairs bacterial clearance while enhancing tissue injury.

Previous studies reported the involvement of neutrophil proteases and specifically elastase in various inflammatory states both in animal models (39–41) and in seriously ill patients (39, 40, 42–44). Our study extends these observations, suggesting that a broad spectrum of proteases participate in dysregulation of neutrophil function in sepsis. This may provide an explanation for the inconsistent results of human trials with elastase-inhibitors (45, 46). Our study did not assess the overall contribution of neutrophil dysfunction induced by circulating peptides to the complex pathophysiology of sepsis. Our results call attention to the potential role of proteolytic degradation products in the pathogenesis of sepsis that warrant further study. Disruption of this feedback loop may be a therapeutic target to improve outcomes in septic patients.

## Materials and Methods

### Materials

Saponin was from Merck (Darmstadt, Germany); cytochrome c (C7752-5g), phorbolmyristate acetate (PMA) (P8139), propidium iodide (p4170), annexin V-FITC (APOAF), lipase (L3126-25G), DN-ase I (D 5025), superoxid-dismutase (SOD) (S2515), lysozyme (L6751), formyl-methionyl-leucyl-phenylalanin (fMLF) (47729), cytochalasin B (c8273), ficoll (17-1440-02), and antibody CD63-PE (Sigma-Aldrich Cat# SAB4700218, RRID:AB_10898448) were from Sigma (St. Louis, MO, USA); sterile endotoxin-free HBSS (137 mM NaCl, 5.4 mM KCl, 0.25 mM Na2HPO4, 5.55 mM glucose, 0.44 mM KH2PO4, 1.3 mM CaCl2, 1.0 mM MgSO4, 4.2 mM NaHCO3 (sh30268.02)), antibodies against CD35 (CD35-FITC; Thermo Fisher Scientific Cat# 11-0359-42, RRID:AB_11042563), and CD66b (CD66b-FITC; Catalog # (Thermo Fisher Scientific Cat# 11-0666-42, RRID:AB_2572461) from ThermoScientific (Waltham, MA, USA); Proteinase K (3115879001) was from Roche (Basel, Switzerland). CM-H2DCFDA General Oxidative Stress Indicator (Cat. No: C6827, also called as “C400”) was from Invitrogen-Thermo Fischer Scientific (MA, USA). LDH test kit (ab65393) was from Abcam (CT, USA). Neutrophil Elastase Colorimetric Drug Discovery Kit (NECDC Kit, BML-AK497-0001) was from Enzo (NY, USA). All other reagents were of research grade. GFP-expressing S. aureus (47) was a kind gift of Professor William Nauseef (University of Iowa, IA, USA). ATCC No. of Meticillin Sensitive *S. aureus* (MSSA), and of *E. coli* is 29213 and ML-35, respectively.

### Patients and healthy volunteers

The study consists of 31 patients admitted to the Clinic for Anesthesiology and Intensive Care of Semmelweis University and of 16 healthy volunteers. Both patients and healthy volunteers were 18 years of age or older and provided informed consent as approved by the Institutional Review Board of Semmelweis University. Inclusion criteria of patients were: clinical proof or strong indication of abdominal or pulmonary infection with onset no longer than 2 days prior to entry plus at least one symptom in each of the general, inflammatory and hemodynamic categories of the diagnostic criteria of sepsis for adult patients, as recommended by the SCCM/ESICM/ACCP/ATS/SIS sepsis consensus conference (7). Immunosuppressive treatment, acute exacerbation of severe chronic illness, polytrauma, known neoplastic disease or disagreement were exclusion criteria. Samples were taken on the 1^st^ (29 samples, APACHE II range: 10-34, average 20.72 ± 6.18 SEM), 7^th^ (25 samples, APACHE II range: 8-32, average 17.04 ± 5.41 SED), and on the 30^th^ (3 samples, APACHE II range: 10-14, average 12.33 ± 2.08 SED) day of treatment. APACHE II score that summarizes the clinical status of the patients on the basis of 12 parameters (48) was calculated at each sampling with Medscape online calculator (http://reference.medscape.com/calculator/apache-ii-scoring-system). The detailed patient data are summarized in Table 1. The total mortality rate among patients up to day 30 was 45.16 %.

**Table 1.** Summary of the clinical parameters of enrolled patients and healthy donors.

|  |  | SepticPatients |  | Healthydonors |  |
| --- | --- | --- | --- | --- | --- |
| Gender | Male | 58,06% | 18 / 31 | 57,14% | 8 / 14 |
|  | Female | 41,94% | 13 / 31 | 42,86% | 6 / 14 |
| Age | Mean (+/- SD) | 70,03 | 12,156 | 59,21 | 6,435 |
|  | Meadian (range) | 71,00 | (41-89) | 60 | (53-71) |
| Infectiosus agent | Gram + | 32,26% | 9 / 31 |  |  |
|  | Gram - | 19,35% | 6 / 31 |  |  |
|  | Fungi | 12,90% | 4 / 31 |  |  |
|  | unknown | 38,71% | 12 / 31 |  |  |
| Localisation of infection | Pulmonal | 54,84% | 17 / 31 |  |  |
|  | Abdominal | 48,39% | 15 / 31 |  |  |
| Anamnesitic data | Chronic obstructive Pulmonal Disease | 16,13% | 5 / 31 | 14,29% | 2 / 14 |
|  | Ischaemic Heart Disease | 38,71% | 12 / 31 | 35,71% | 5 / 14 |
|  | Diabetes Mellitus (Type I or II) | 48,39% | 15 / 31 | 42,86% | 6 / 14 |
|  | Stroke | 6,45% | 2 / 31 | 7,14% | 1 / 14 |
|  | Hypertonia | 74,19% | 23 / 31 | 64,29% | 9 / 14 |
|  | Chronic Renal Diseases | 0% | 0 / 31 | 0% | 0 / 14 |
|  | Hepatopathia chronica | 3,23% | 1 / 31 | 0% | 0 / 14 |
|  | Immundeficiency | 0% | 0 / 31 | 0% | 0 / 14 |
|  | BMI Mean (+/- SD) | 28,9642 | 4,270 | 26,186 | 6,435 |
|  | Meadian (range) | 29,3848 | (20.2-36.7) | 28,0 | (23.4-36.5) |
|  | Alcoholol | 19,35% | 6 / 31 | 7,14% | 1 / 14 |
|  | Nicotin | 38,71% | 12 / 31 | 35,71% | 5 / 14 |
| Clinical status | APACHE II score Mean (+/- SD) | 18,57 | 6,198 |  |  |
|  | Meadian (range) | 17,00 | (8-29) |  |  |
|  | APACHE II score of deceased |  |  |  |  |
|  | Mean (+/- SD) | 22,722 | 1,522 |  |  |
|  | Meadian (range) | 24,5 | (12-34) |  |  |
|  | APACHE II score of survivals |  |  |  |  |
|  | Mean (+/- SD) | 16,639 | 0,907 |  |  |
|  | Meadian (range) | 17 | (8-29) |  |  |
|  | Deceased patients | 45,16% | 14 / 31 | 0% | 0 / 14 |
|  | Survival patients | 54,84% | 17 / 31 | 100% | 14 / 14 |

### Preparation of PMN

Venous blood was drawn into citrate tubes from adult septic patients and healthy volunteers according to procedures approved by the Institutional Review Board of the Semmelweis University. Neutrophils were obtained by dextran sedimentation followed by Ficoll-Paque gradient centrifugation as described previously (34). The preparation contained over 95% PMN and less than 0.5% eosinophils. Preparations were checked by May-Grünwald-Giemsa stain of sample smears.

### Preparation of monocytes, lymphocytes, red blood cells and blood plasma

Human Peripheral Blood Mononuclear Cell (PBMC) fraction was separated by Ficoll-Paque gradient centrifugation, as supernatant fraction of PMN sedimentation. Monocytes were separated by RosetteSep Human Monocyte Enrichment cocktail, following the manufacturer’s instructions (Stemcell Technologies, Vancouver, Canada). Lymphocytes were separated by RosetteSep Human T cell Enrichment and by RosetteSep Human B cell Enrichment cocktail, following the manufacturer’s instructions (Stemcell Technologies). Preparations were checked by May-Grünwald-Giemsa stain of smears of samples. For preparation of red blood cells, 1 ml of citrated blood of healthy donor was sedimented with 500 x g for 1 minute. After removal of supernatant, the sediment was resuspended in 1ml PBS. Density of cells was determined in Bürker chamber after 10^4^ times dilution. Blood plasma was separated by centrifugation of samples (5000 x g, 5 min, room temperature). Blood plasma samples for further investigations were stored at −80 °C.

### Opsonization of bacteria

Bacteria (10^9^/ml in HBSS) were opsonized with 10 % (v/v) pooled normal human serum (acquired at least from 4 healthy donors) for 20 minutes at 37°C. After opsonization, bacteria were sedimented (5 minutes, 4°C, 5000 x g), and washed with HBSS. The concentration of bacteria was set to 10^9^/ml on the basis of optical density (OD600 = 1.0).

### Measurement of bacterial survival

Indicated amount of opsonized bacteria (suspended in 100 μl HBSS) were incubated with 900 μl, 10^7^/ml PMN (diluted in HBSS) for 30 minutes with gentle (160 / min.) shaking. Samples (100 μl) were transferred at the beginning and every 10 minutes to ice-cold HBSS, containing 1mg/ml saponin and kept at −80°C for 20 minutes to disrupt PMN. Freezing and saponin dissolved in HBSS did not impair subsequent growth of the used bacteria. Original CFUs were estimated with calibration samples of the applied bacteria, based on changes in optical density at 650 nm. Growth of bacteria was followed with ELISA plate reader (Labsystems iEMS Reader MF, Thermos Scientific), as previously described (34).

### Measurement of Reactive Oxygen Species

Release of extracellular superoxide (O_2_^.-^) was determined with superoxide dismutase (SOD) inhibitable cytochrome C reduction system, as previously described (34). The SOD-insensitive value was subtracted from all samples.

Intracellular reactive oxygen species production was measured either by lucigenin-based chemiluminescent method, (as previously described (34), using a Mithras LB 940 multilabel reader (Berthold, Germany) or by CM-H2DCFDA General Oxidative Stress Indicator (C400) in confocal fluorescent microscopy (microscope and objectives are described below). For lucigenin-based measurements, parallel samples were measured by cytochrome C reduction assay, and samples were correlated on the basis of non-activated and 100 nM PMA activated samples. Values of non-activated samples were subtracted. As bacterial stimuli, 10^8^/ml opsonized *S. aureus* or *E. coli* was used. For C400 measurements one aliquot (50 μg) of C400 was dissolved in 125 μl of DMSO. 10^7^ / ml PMN in PBS were incubated with 100 times diluted C400 solution (final concentration: 4 μg/ml) for 10 min at 37 °C with gentle shake, followed by centrifugation (2000 x g 30 sec, room temperature), and the sediment was taken up in fresh (HBSS or PBS) solution. Samples were incubated in microscopic chambers (see at microscopy) at 37 °C with 3×10^9^/ml opsonized *S. aureus* for 20 minutes. Background fluorescence intensity was subtracted from all samples. Fluorescence intensity of each sample was normalized to fluorescence intensity of GFP expressing *S. aureus* (USA 300). For controlling functional integrity of PMN after the staining procedure with C400, fluorescence intensity of untreated and 100 nM PMA treated PMN was followed in time.

### Confocal video microscopy

For confocal video microscopy experiments Zeiss LSM710 confocal laser scanning microscope equipped with 40 ×/1.3 water objective (Plan-Neofluar, Zeiss) was used, with 1μm optical slices. In live/video microscopic experiments stage and objective were heated to 37°C and microscopic slips were coated prior to use with heat inactivated BSA (10 wt/vol %) for 1 hour at room temperature and were washed with HBSS two times. Live measurements were carried out in heated microscopic chambers in HBSS solutions. For measurement of C400 intensity, 488 nm wavelength was used for excitation, while emission was detected in the range of 500 and 550 nm. For appropriate parallel distinguishing of C400 and GFP, all images were captured in multitrack mode with ZEN 2.3 software (Zeiss). In these cases 488 nm wavelength was used for excitation, while detection of emission with multitrack started at 490 nm, with 32 tracks, with 9 nm length of each tracks, up to 778 nm. Successful distinguishing of C400 and GFP was controlled by only C400 filled cells and by only GFP containing bacteria samples. In all cases non-stained cells and GFP expressing bacteria were used for proper adjustment of detection. Images were analyzed with ZEN 2.3 software (Zeiss) or by Image J 1.42 software (NIH). For colocalization measurements ZEN 2.3 software (Zeiss) was used, where intensity limits for colocalization were selected on the basic on non-stained samples.

### Phagocytic activity of PMN

PMN (4.5 x 10^6^ in 450 μl HBSS) were incubated with two different amounts (10^8^, or 3×10^9^ CFU / ml final concentration) of opsonized GFP-expressing *S. aureus* (USA300) for 20 minutes. PMN were identified on the basis of FSC and SSC by flow cytometry. Amount of phagocytic PMN was determined in a Becton Dickinson FACSCalibur Analytic Flow cytometer based on GFP-positivity. Phagocytic index (number of phagocytosed bacteria / cell) was determined in parallel calibration samples, analyzed by confocal microscopy.

### Measurement of PMN degranulation

PMN (10^5^ naïve cells, suspended in 300 μl PBS) were incubated with 300 μl HBSS (negative control), or with 300 μl HBSS with 1 μM of fMLP and 10 μM cytochalasin B (positive control) or with 300 μl of 3-12 kDa fraction of marked sample for 1 hour at 37 °C with gentle shake. After incubation all samples were sedimented by 2000g 30 sec room temperature centrifugation and PMN were resuspended in 100 μl fresh HBSS. Samples (90 μl) were stained with anti-CD35-FITC or with anti-CD63-PE or with anti-CD66b-FITC primary conjugated antibodies, following the manufacturer’s protocol. Non-stained naïve PMN were used as negative staining control. Fluorescence was measured by BD FacsCalibur flow cytometer. PMN were identified on the basis of forward and side scatter, fluorescence intensity of PMN was compared to the non-stained sample. The values for the negative and positive controls are shown in Fig. S3e.

### Treatment of blood plasma

For heat inactivation, plasma was treated at 100 °C for 15 minutes with shaking (600 rpm). After heat treatment, samples were centrifuged at 22000 x g for 20 minutes at 4 °C and the supernatant was filtered through Millex GV 220 nm filter (Merck-Millipore). For acid treatment acetic acid was used with 0.1 N final concentration, for 20 minutes at room temperature. pH was restored to 7.4 by slow mixing of NaOH. Dialysis of 1 ml plasma against a final volume of 25 ml PBS was carried out for 24 hours at 4 °C using Membra-Cel MC24 (Sigma St. Louis, MO, USA) with gentle shake. Membrane cut-off was indicated by the producer to be approx.10 kDa. We tested that cytochrome c (MW: ~12 kDa) was fully retained. Dialyzing conditions were established by dialysis of Trypane blue (MW: 0.96 kDa) and protein concentration measurement series in the external dialyzing fluid during dialysis of proteinase K digested blood plasma from healthy donors (plateau was reached at 20 hour of dialysis, data not shown). External solution was changed 4 times during dialysis. After dialysis external solutions were mixed and were concentrated by Centricon YM-3 (from Merck-Millipore, used at 5000 x g for 30 minutes, 4 °C) to reach 1 ml as final volume. For protein degradation of plasma, 200 μg/ml Proteinase K was used for 60 minutes at 37 °C. For lipid degradation, 75 U / ml (measured against triacetin) porcine pancreas lipase was used for 60 minutes at 37 °C. For DNA elimination 1 U/μl DNase I was used for 30 minutes at room temperature. For carbohydrate degradation 0.5 μg/ml lysozyme was used at 37 °C for 30 minutes. All enzymatic reactions were stopped with heat treatment (100 °C, 15 min.) of samples. Protein concentration was determined by the Bradford protein assay using BSA as standard.

### Production and treatment of cell lysates

10^7^ WBC, 10^8^ RBC or 10^9^ bacteria (diluted in 1 ml HBSS) were centrifuged (1000 x g for 5 min for cells, 5000 x g for 5 min for bacteria, room temperature), followed by snap-freezing with liquid nitrogen, and lysing by ultrasonic treatment with 5 x 10 sec cycles. Lysis of cells was checked in a Becton Dickinson FACSCalibur Analytic Flow cytometer. This procedure was repeated until detectable cell or bacteria count was less than 1 % of initial cell count. Where marked, cell lysates were treated either with 10 times diluted PMN lysate (for 1 hour) or by 20 μg/ml Proteinase K (for 10 min.). Each sample was treated with heat and dialyzed as described above.

### Treatment of PMN by blood plasma or by cell lysates

Typically, 2 ml of blood plasma derived samples or cell lysate samples were incubated with 2 ml of 10^7^ / ml PMN in PBS for 60 min with gentle shake at 37 °C, followed by centrifugation (2000 x g, 30 sec, room temperature). Cells were resuspended in fresh HBSS and their count and viability was controlled on the basis of propidium iodide and Annexin V-FITC binding to cells in Becton Dickinson FACSCalibur Analytic Flow cytometer, following the manufacturer’s protocol. As positive control, PMA treated (100 nM, 2h, 37 °C, gentle shake) PMN were used.

### Measurement of neutrophil elastase (NE) and total protease activity of blood plasma

For NE activity measurement, Neutrophil Elastase Colometric Drug Discovery Kit (NECDC Kit) (Enzo) was modified as follows: 65 μl of NECDC Kit buffer with 5 μl of BML-P231-9090 NE substrate was mixed with 30 μl of blood plasma, and was incubated in ELISA plate reader (Labsystems iEMS Reader MF, Thermos Scientific) at 37 °C with gentle shake for 2 hours, with measuring OD at 405 nm at every minute. Each sample had a parallel, co-incubated with 4 mM BML-PI103-9090 (Elastatinal) as NE inhibitor. Values of NE inhibited samples were subtracted from non-inhibited sample values. Neutrophil Elastase was used as positive control, alongside with an inhibitory parallel. The results were analyzed at saturation of the reaction.

For protease activity measurement of blood plasma, 30 μl of 2 mg/ml Ovalbumin (OVA) (Sigma) was incubated for 12 h at 37 °C with gentle shake with 5 μl of blood plasma in 25 μl of PBS in the presence or absence of 1% (w/v) of Protease Inhibitor Cocktail from Sigma (PIC). As control, lysate of PMN (5 μl of a 10^7^ PMN / ml solution) was used. Samples were run on SDS-PAGE and stained by Coomasie G250. OVA concentration was assessed on the basis of calibration curve in the concentration range from 0.03 mg/ml to 3 mg/ml). For determining Coomasie intensity, Image J 1.42 software (NIH) was used.

### Co-incubation of PBMC, PMN or RBC with plasma-fraction treated PMN

PBMC, PMN and RBC were isolated as described above from healthy donors. PMN (10^7^ PMN in 1 ml PBS) from the identical donor were incubated with the activating plasma fraction as described above. An aliquot of 2.7 ml of the indicated target cell (10^7^ / ml for PBMC and PMN, 10^9^ / ml for RBC, all in HBSS) were incubated with 0.3 ml of 10^7^ / ml activating-fraction treated PMN for 1 hour at 37 °C with gentle shake, followed by centrifugation (500 x g for 5 min, room temperature). The target cells were examined by AnnexinV-Propidium iodide (PBMC, PMN) or by Erytrosin B (RBC) assay, following the manufacturer’s protocol, in Becton Dickinson FACSCalibur Analytic Flow cytometer. Cells were identified on the basis of forward- and side scatter. Supernatants were examined either with LDH assay (following the manufacturer’s protocol) or by Hemoglobin release assay.

### Hemoglobin (HGB) release assay

2 x 1 ml of 10^9^ / ml RBC (in HBSS) from the same donor were sedimented (500 x g for 5 min, room temperature). One aliquot was re-suspended in distilled water (maximal hemolysis), the other one was re-suspended in HBSS as negative control. Optical density of supernatant of the same amount of samples (as described above) were compared to the OD of these 2 samples at 405 nm.

### LC-MS analysis of plasma protein degradome

#### Data Collection

Samples were diluted to a final concentration of 0.08μg/μL in 2% v/v acetonitrile/0.4% v/v formic acid. Samples were injected on to a μ-Precolumn^™^ Cartridge (Acclaim PepMap100 C18, 5 μm, 100 Å, 300 μm ID x 5 mm from Thermo Fisher Scientific) and peptides resolved on an 360 μm OD x 100 μm ID fused silica tip packed with 12cm of Aeris Peptide 3.6 μm XB-C18 100Å material (Phenomenex, Torrance, CA, USA). An EASY-nLC 1000 System was used to elute peptides under buffer A (2% v/v acetonitrile/0.1% v/v formic acid) to buffer B (80% v/v acetonitrile/0.1% v/v formic acid) at 300 nL/min with a 100 min linear gradient from 0% B to 50% B. Eluted peptides were introduced by nanoelectrospray (Nanospray Flex source (Thermo Fisher) at a 225°C ion transfer capillary temperature and spray voltage of 1.6kV into an Orbitrap Elite–ETD mass spectrometer (Thermo Fisher). Data was collected using an Nth Order Double Play with ETD Decision Tree method created in Xcalibur v2.2 (Thermo Fisher). Scan event one obtained an FTMS MS1 scan (normal mass range; 60,000 resolution, full scan type, positive polarity, profile data type) for the range 300-2000 m/z. Scan event two obtained ITMS MS2 scans (normal mass range, rapid scan rate, centroid data type) on up to twenty peaks that had a minimum signal threshold of 10,000 counts from scan event one. A decision tree was used to determine whether CID or ETD activation was used. An ETD scan was triggered if any of the following held: an ion had charge state 3 and m/z less than 650, an ion had charge state 4 and m/z less than 900, an ion had charge state 5 and m/z less than 950, or an ion had charge state greater than 5; a CID scan was triggered in all other cases. The lock mass option was enabled (0% lock mass abundance) using the 371.101236 m/z polysiloxane peak as an internal calibrant.

#### Data Analysis

RAW files were analyzed in Peaks Studio 7.5 (Bioinformatics Solutions Inc., Waterloo, ON, Canada) using the Denovo, PeaksDB, PeaksPTM, and Label Free Q algorithms. The 7/14/2016 version of the UniprotKB reference proteome canonical and isoform *Homo sapiens* sequences (Proteome ID UP000005640) concatenated with the gpm.org cRAP database. No enzyme was specified in the search with precursor and fragment tolerances of 50 ppm and 1.0 Da, respectively. Feature.csv data were exported from the Peaks Studio 7.5 Label Free Q result. Total ion chromatogram (TIC) normalized peak intensities were exported into excel and formatted for statistical analysis.

### Calculations based on proteomics analysis

To predict the amount of a given peptide in a given sample, first the relative amount of all peptides was determined, on the basis of reference average area of the given peptide and the relative abundance in the given sample (relative amount of a peptide = ref. Avg. Area x relative abundance). These relative amounts were summarized for all the peptides in the given sample and were normalized to the total amount of peptide in the sample (measured by Bradford method).

For prediction of intrinsic disorder in the structure of the peptide OneDCRF (Uni. Umea, Sweden) online program(49) (http://babel.ucmp.umu.se/ond-crf/) and ModiDB-lite(50) (of University of Padova) (http://protein.bio.unipd.it/mobidblite/) were used, following the instructions of the manufacturers. Only those peptides are presented where these 2 programs provided identical results.

To predict the protease enzymes that could be responsible for generation of peptides, Proteasix online program, Prediction Tool, Observed mode(51) was used (http://www.proteasix.org/), following the manufacturer’s instructions. To determine the importance of an enzyme in generation of the peptides of a given sample, the amount of all peptides produced by the given enzyme was summarized.

### Statistical analysis

Results are presented as mean (+/- SEM) of indicated number of independent experiments. Data were compared with Mann-Whitney u-test, or with Student’s t-test, or with one-way ANOVA test followed by a Tukey post hoc test, depending on the condition, using Statistica 11 software (StatSoft). Correlations also were investigated with Statistica 11 software after testing normal distribution of the data (StatSoft).

### Study approval

Written informed consent was received from participants prior to inclusion in the study as approved by the Institutional Review Board of Semmelweis University (for septic patients: 69/2009 to Zs.I., for healthy donors: 40/1998, 021/01563-2/2015 to E.L.)

## Authorship contributions

Cs.I.T. designed and carried out the majority of experiments, summarized data, prepared the figures and participated in writing of the manuscript.

F.K. carried out the experiments on neutrophil function under different environmental conditions and helped Cs.I.T. in many experiments.

V.B. and E.T. selected the patients and monitored the clinical parameters. V.B. also helped in establishing the protocol for plasma treatment of PMN.

A.P. carried out the majority of experiments on cell lysates.

K.R.M., M.L.M. and D.W.W. carried out the proteomic analysis, K.R.M. helpedediting the manuscript.

Zs.I. initiated the study and obtained the permission from the Institutional Review Board. E.L. directed, oversaw and obtained financing for the experiments, discussed the results and prepared the manuscript.

## Acknowledgements

The authors are indebted to Professor Péter Tompa, Drs. Gabriella Pócsfalvi and Gitta Strasser for stimulating discussions, to Professor Attila Mócsai, Drs. Ákos M. Lőrincz and Roland Csépányi-Kömi for critical reading of the manuscript and Ms. Regina Toth-Kun and Ms. Edit Fedina for devoted and excellent technical assistance. Experimental research was supported by grants from the Hungarian Research Fund (OTKA K108382, K119236) and from VEKOP project (2.3.2-0016) to E.L.

## References

1. Singer M, Deutschman CS, Seymour CW, Shankar-Hari M, Annane D, Bauer M, et al. The Third International Consensus Definitions for Sepsis and Septic Shock (Sepsis-3). JAMA. 2016;315(8):801–10.

2. Annane D, Bellissant E, and Cavaillon JM. Septic shock. Lancet. 2005;365(9453):63–78.

3. Brown KA, Brain SD, Pearson JD, Edgeworth JD, Lewis SM, and Treacher DF. Neutrophils in development of multiple organ failure in sepsis. Lancet. 2006;368(9530):157–69.

4. Namas R, Zamora R, An G, Doyle J, Dick TE, Jacono FJ, et al. Sepsis: Something old, something new, and a systems view. J Crit Care. 2012;27(3):314 e1–11.

5. Gaieski DF, Edwards JM, Kallan MJ, and Carr BG. Benchmarking the incidence and mortality of severe sepsis in the United States. Critical care medicine. 2013;41(5):1167–74.

6. Cohen J. The immunopathogenesis of sepsis. Nature. 2002;420(6917):885–91.

7. Levy MM, Fink MP, Marshall JC, Abraham E, Angus D, Cook D, et al. 2001 SCCM/ESICM/ACCP/ATS/SIS International Sepsis Definitions Conference. Critical care medicine. 2003;31(4):1250–6.

8. Vincent JL, Sakr Y, Sprung CL, Ranieri VM, Reinhart K, Gerlach H, et al. Sepsis in European intensive care units: results of the SOAP study. Critical care medicine. 2006;34(2):344–53.

9. Zahar JR, Timsit JF, Garrouste-Orgeas M, Francais A, Vesin A, Descorps-Declere A, et al. Outcomes in severe sepsis and patients with septic shock: pathogen species and infection sites are not associated with mortality. Critical care medicine. 2011;39(8):1886–95.

10. Treacher DF, Sabbato M, Brown KA, and Gant V. The effects of leucodepletion in patients who develop the systemic inflammatory response syndrome following cardiopulmonary bypass. Perfusion. 2001;16 Suppl:67–73.

11. El Kebir D, Gjorstrup P, and Filep JG. Resolvin E1 promotes phagocytosis-induced neutrophil apoptosis and accelerates resolution of pulmonary inflammation. Proc Natl Acad Sci U S A. 2012;109(37):14983–8.

12. McGill SN, Ahmed NA, Hu F, Michel RP, and Christou NV. Shedding of L-selectin as a mechanism for reduced polymorphonuclear neutrophil exudation in patients with the systemic inflammatory response syndrome. Arch Surg. 1996;131(11):1141–6; discussion 7.

13. Tavares-Murta BM, Zaparoli M, Ferreira RB, Silva-Vergara ML, Oliveira CH, Murta EF, et al. Failure of neutrophil chemotactic function in septic patients. Critical care medicine. 2002;30(5):1056–61.

14. Reddy RC, Narala VR, Keshamouni VG, Milam JE, Newstead MW, and Standiford TJ. Sepsis-induced inhibition of neutrophil chemotaxis is mediated by activation of peroxisome proliferator-activated receptor-{gamma}. Blood. 2008;112(10):4250–8.

15. Chishti AD, Shenton BK, Kirby JA, and Baudouin SV. Neutrophil chemotaxis and receptor expression in clinical septic shock. Intensive Care Med. 2004;30(4):605–11.

16. Martins PS, Brunialti MK, Martos LS, Machado FR, Assuncao MS, Blecher S, et al. Expression of cell surface receptors and oxidative metabolism modulation in the clinical continuum of sepsis. Crit Care. 2008;12(1):R25.

17. El Kebir D, and Filep JG. Modulation of Neutrophil Apoptosis and the Resolution of Inflammation through beta2 Integrins. Front Immunol. 2013;4:60.

18. Kennedy AD, and DeLeo FR. Neutrophil apoptosis and the resolution of infection. Immunol Res. 2009;43(1-3):25–61.

19. Geering B, Stoeckle C, Conus S, and Simon HU. Living and dying for inflammation: neutrophils, eosinophils, basophils. Trends Immunol. 2013;34(8):398–409.

20. Simms HH, and D’Amico R. Polymorphonuclear leukocyte dysregulation during the systemic inflammatory response syndrome. Blood. 1994;83(5):1398–407.

21. Martins PS, Kallas EG, Neto MC, Dalboni MA, Blecher S, and Salomao R. Upregulation of reactive oxygen species generation and phagocytosis, and increased apoptosis in human neutrophils during severe sepsis and septic shock. Shock. 2003;20(3):208–12.

22. Kaufmann I, Hoelzl A, Schliephake F, Hummel T, Chouker A, Peter K, et al. Polymorphonuclear leukocyte dysfunction syndrome in patients with increasing sepsis severity. Shock. 2006;26(3):254–61.

23. Wenisch C, Parschalk B, Patruta S, Brustbauer R, and Graninger W. Effect of polyclonal immunoglobulins on neutrophil phagocytic capacity and reactive oxygen production in patients with gram-negative septicemia. Infection. 1999;27(3):183–6.

24. Fung YL, Fraser JF, Wood P, Minchinton RM, and Silliman CC. The systemic inflammatory response syndrome induces functional changes and relative hyporesponsiveness in neutrophils. J Crit Care. 2008;23(4):542–9.

25. Raymond SL, Hawkins RB, Stortz JA, Murphy TJ, Ungaro R, Dirain ML, et al. Sepsis is associated with reduced spontaneous neutrophil migration velocity in human adults. PLoS One. 2018;13(10):e0205327.

26. Timar CI, Lorincz AM, Csepanyi-Komi R, Valyi-Nagy A, Nagy G, Buzas EI, et al. Antibacterial effect of microvesicles released from human neutrophilic granulocytes. Blood. 2013;121(3):510–8.

27. Shi Z, Chen K, Liu Z, Sosnick TR, and Kallenbach NR. PII structure in the model peptides for unfolded proteins: studies on ubiquitin fragments and several alanine-rich peptides containing QQQ, SSS, FFF, and VVV. Proteins. 2006;63(2):312–21.

28. Tompa P, and Csermely P. The role of structural disorder in the function of RNA and protein chaperones. FASEB J. 2004;18(11):1169–75.

29. Whittington SJ, Chellgren BW, Hermann VM, and Creamer TP. Urea promotes polyproline II helix formation: implications for protein denatured states. Biochemistry. 2005;44(16):6269–75.

30. Hackl EV. Effect of temperature on the conformation of natively unfolded protein 4E-BP1 in aqueous and mixed solutions containing trifluoroethanol and hexafluoroisopropanol. Protein J. 2015;34(1):18–28.

31. Sengelov H, Kjeldsen L, and Borregaard N. Control of exocytosis in early neutrophil activation. J Immunol. 1993;150(4):1535–43.

32. Sengelov H, Follin P, Kjeldsen L, Lollike K, Dahlgren C, and Borregaard N. Mobilization of granules and secretory vesicles during in vivo exudation of human neutrophils. J Immunol. 1995;154(8):4157–65.

33. McLeish KR, Merchant ML, Creed TM, Tandon S, Barati MT, Uriarte SM, et al. Frontline Science: Tumor necrosis factor-alpha stimulation and priming of human neutrophil granule exocytosis. J Leukoc Biol. 2017;102(1):19–29.

34. Rada BK, Geiszt M, Kaldi K, Timar C, and Ligeti E. Dual role of phagocytic NADPH oxidase in bacterial killing. Blood. 2004;104(9):2947–53.

35. van der Lee R, Buljan M, Lang B, Weatheritt RJ, Daughdrill GW, Dunker AK, et al. Classification of intrinsically disordered regions and proteins. Chem Rev. 2014;114(13):6589–631.

36. Uriarte SM, Rane MJ, Luerman GC, Barati MT, Ward RA, Nauseef WM, et al. Granule exocytosis contributes to priming and activation of the human neutrophil respiratory burst. J Immunol. 2011;187(1):391–400.

37. Nauseef WM. Myeloperoxidase in human neutrophil host defence. Cell Microbiol. 2014;16(8):1146–55.

38. Nauseef WM. How human neutrophils kill and degrade microbes: an integrated view. Immunol Rev. 2007;219:88–102.

39. Catz SD, and McLeish KR. Therapeutic targeting of neutrophil exocytosis. J Leukoc Biol. 2020;107(3):393–408.

40. Potey PM, Rossi AG, Lucas CD, and Dorward DA. Neutrophils in the initiation and resolution of acute pulmonary inflammation: understanding biological function and therapeutic potential. J Pathol. 2019;247(5):672–85.

41. Altshuler AE, Penn AH, Yang JA, Kim GR, and Schmid-Schonbein GW. Protease activity increases in plasma, peritoneal fluid, and vital organs after hemorrhagic shock in rats. PLoS One. 2012;7(3):e32672.

42. Bauza-Martinez J, Aletti F, Pinto BB, Ribas V, Odena MA, Diaz R, et al. Proteolysis in septic shock patients: plasma peptidomic patterns are associated with mortality. Br J Anaesth. 2018;121(5):1065–74.

43. Fujishima S, Morisaki H, Ishizaka A, Kotake Y, Miyaki M, Yoh K, et al. Neutrophil elastase and systemic inflammatory response syndrome in the initiation and development of acute lung injury among critically ill patients. Biomed Pharmacother. 2008;62(5):333–8.

44. Wilkinson TS, Conway Morris A, Kefala K, O’Kane CM, Moore NR, Booth NA, et al. Ventilator-associated pneumonia is characterized by excessive release of neutrophil proteases in the lung. Chest. 2012;142(6):1425–32.

45. Zeiher BG, Artigas A, Vincent JL, Dmitrienko A, Jackson K, Thompson BT, et al. Neutrophil elastase inhibition in acute lung injury: results of the STRIVE study. Critical care medicine. 2004;32(8):1695–702.

46. Polverino E, Rosales-Mayor E, Dale GE, Dembowsky K, and Torres A. The Role of Neutrophil Elastase Inhibitors in Lung Diseases. Chest. 2017;152(2):249–62.

47. Schwartz J, Leidal KG, Femling JK, Weiss JP, and Nauseef WM. Neutrophil bleaching of GFP-expressing staphylococci: probing the intraphagosomal fate of individual bacteria. Journal of immunology. 2009;183(4):2632–41.

48. Knaus WA, Draper EA, Wagner DP, and Zimmerman JE. APACHE II: a severity of disease classification system. Critical care medicine. 1985;13(10):818–29.

49. Wang L, and Sauer UH. OnD-CRF: predicting order and disorder in proteins using [corrected] conditional random fields. Bioinformatics. 2008;24(11):1401–2.

50. Necci M, Piovesan D, Dosztanyi Z, and Tosatto SCE. MobiDB-lite: fast and highly specific consensus prediction of intrinsic disorder in proteins. Bioinformatics. 2017;33(9):1402–4.

51. Klein J, Eales J, Zurbig P, Vlahou A, Mischak H, and Stevens R. Proteasix: a tool for automated and large-scale prediction of proteases involved in naturally occurring peptide generation. Proteomics. 2013;13(7):1077–82.

