## supplement figures 1-4 for "Disordered peptides impair neutrophil bacterial clearance and enhance tissue damage in septic patients"

Suppl. Fig. 1.

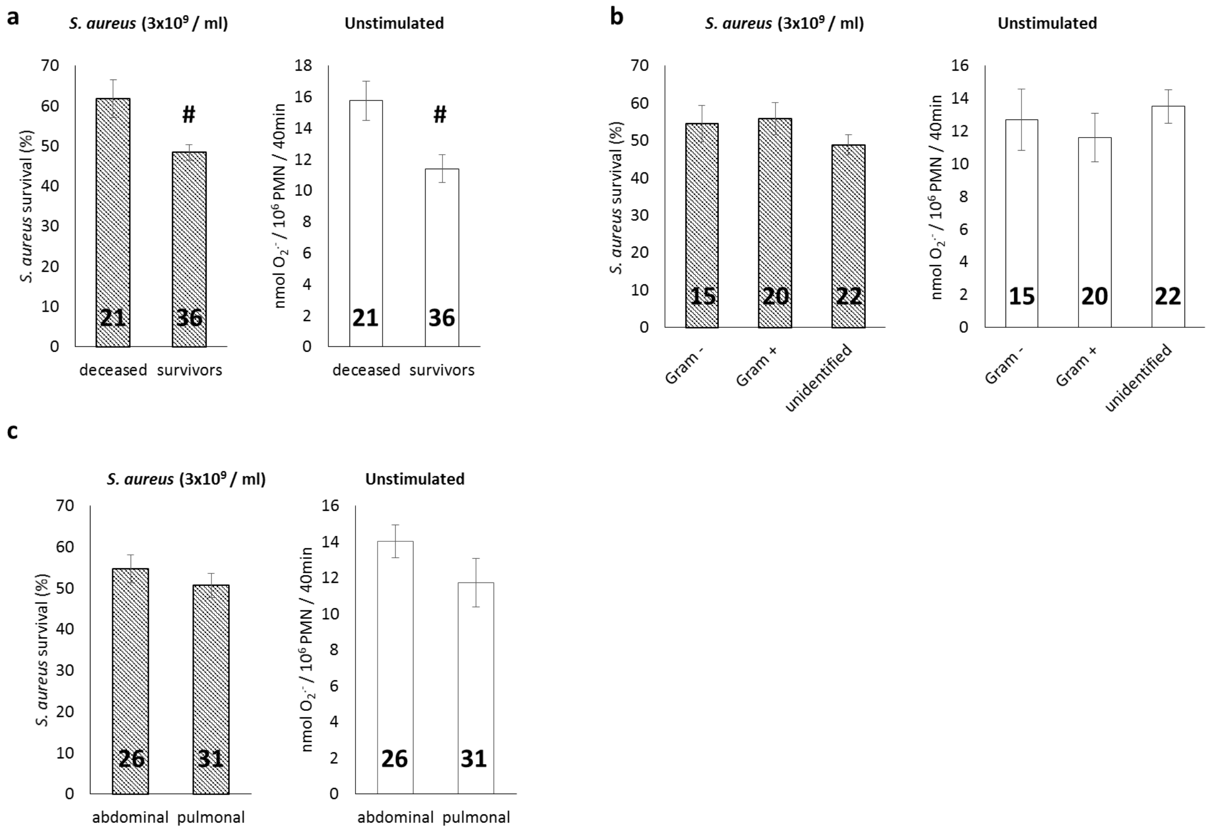


**Analysis of functional properties of PMN from septic patients**.

Patient data were selected according to: **a.:** survival of patients, **b.:** type of clinical microbiologically proved pathogen and **c.:** site of focus of infection. Left panels: survival of $3x10^9$ / ml *S. aureus* during incubation with PMN, right panels: unstimulated superoxide release by PMN. Error bars represents SEM. Indicated numbers in columns represent number of independent samples. #: $p < 0.01$. Significance above $p > 0.05$ is not indicated.

Suppl. Fig. 2.

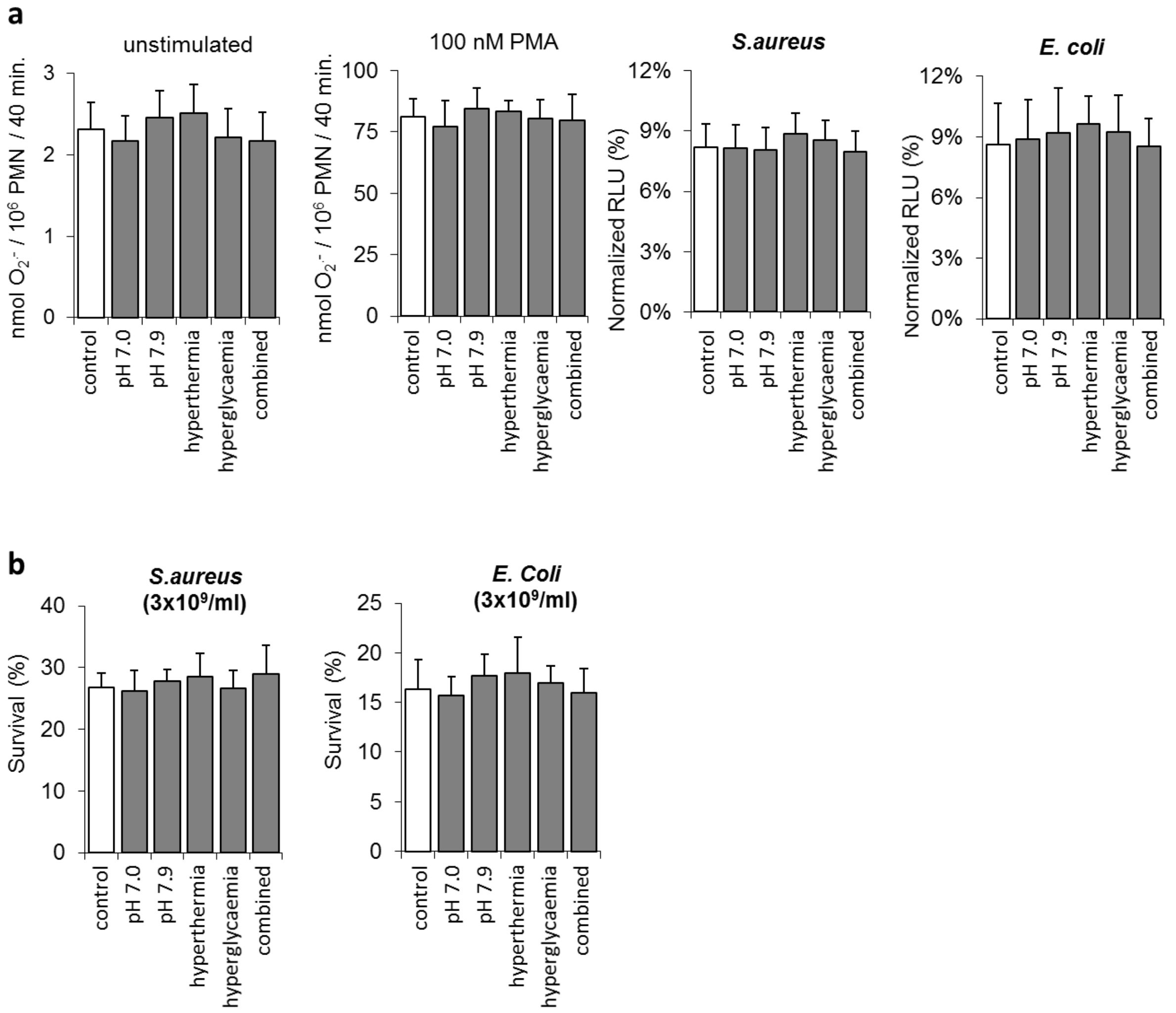


**Functional properties of PMN from healthy donors in different physical and chemical environment during reaction**.

Hyperthermia: 40 °C. Hyperglycemia: glucose concentration: 15mM. Combined: pH=7.0 with hyperthermia (40 °C ) and with hyperglycemia (15 mM glucose). **a.:** unstimulated extracellular superoxide release, 100 nM PMA stimulated extracellular superoxide release, ROS production to phagocytosis of opsonized *S. aureus* or to opsonized *E. coli*, respectively. **b.:** Bacteria survival tests against opsonized *S. aureus* or against *E. coli*, respectively ($3 \times 10^9$ CFU / ml each). ($n = 4$, ±SEM). Significance above $p > 0.05$ is not indicated.

Suppl. Fig. 3.

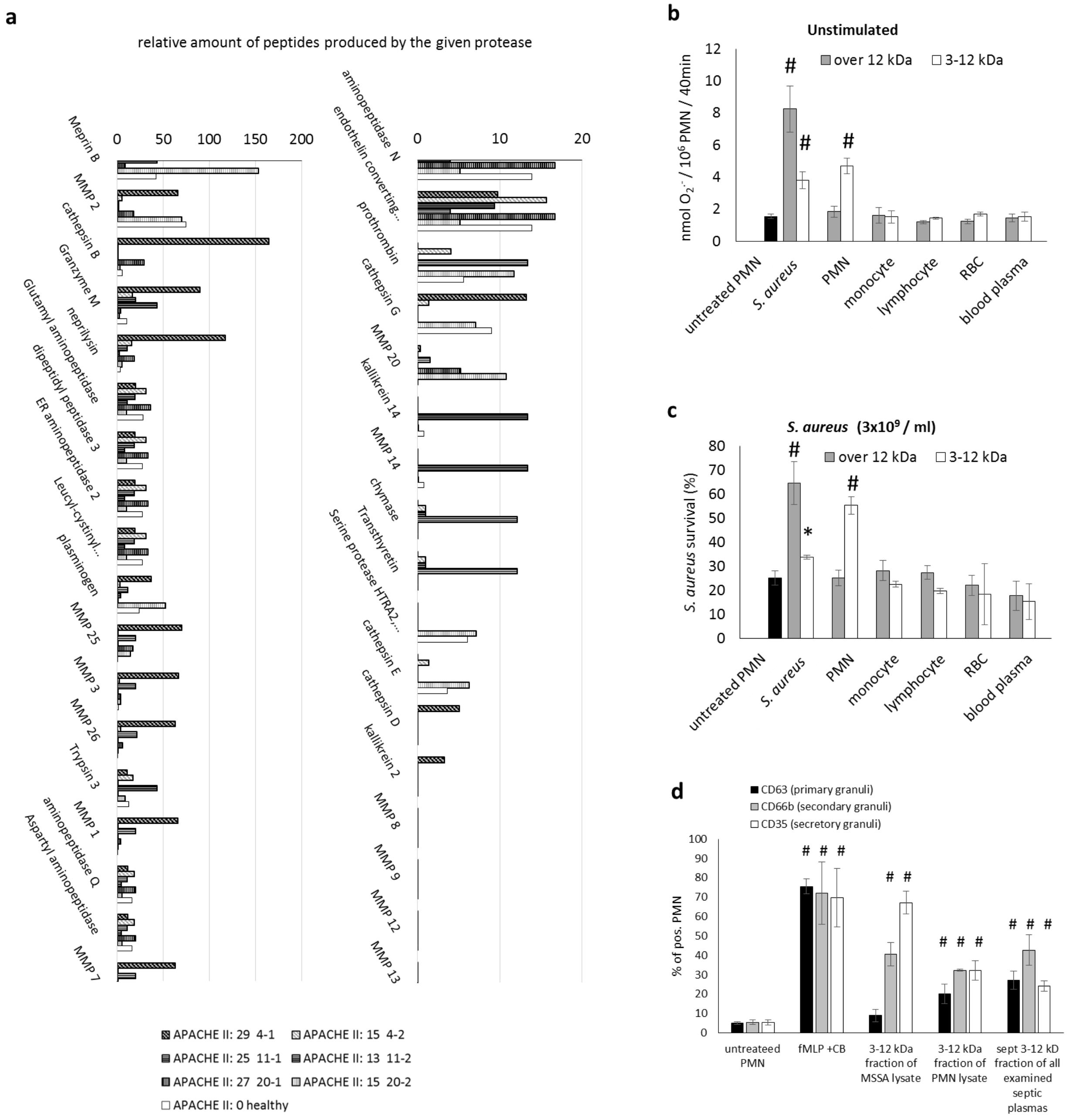


**Potential enzymes and substrates producing 3-12kDa peptides in septic plasma**

**a.:** All proteases, excluded the 5 most common, which can be involved in production of peptides, based on the amount of peptides cleaved by them in the seven samples mentioned above.

**b., c.:** Effect of 3-12 kDa and over 12 kDa fractions of lysates of *S. aureus*, indicated cells or healthy plasma on function of PMN from healthy donors. **b.:** unstimulated superoxide release, **c.:**

*S. aureus* survival ($3x10^9$CFU/ml) (n=6, ±SEM. #: $p < 0.01$, *: $p < 0.05$). **d.::** Degranulation of naïve PMN, followed by CD35 (secretory vesicles), CD66b (secondary granules) and CD63 (primary granules) positivity, after incubation with the marked samples. fMLP (1 μM) + CB (10 μM) served as positive control. SEM. #: $p < 0.05$.
